# Human loss-of-function variants suggest that partial LRRK2 inhibition is a safe therapeutic strategy for Parkinson’s disease

**DOI:** 10.1101/561472

**Authors:** Nicola Whiffin, Irina M. Armean, Aaron Kleinman, Jamie L. Marshall, Eric V. Minikel, Julia K. Goodrich, Nicholas M Quaife, Joanne B Cole, Qingbo Wang, Konrad J. Karczewski, Beryl B. Cummings, Laurent Francioli, Kristen Laricchia, Anna Guan, Babak Alipanahi, Peter Morrison, Marco A.S. Baptista, Kalpana M. Merchant, Genome Aggregation Database Production Team, Genome Aggregation Database Consortium, James S. Ware, Aki S. Havulinna, Bozenna Iliadou, Jung-Jin Lee, Girish N. Nadkarni, Cole Whiteman, the 23andMe Research Team, Mark Daly, Tõnu Esko, Christina Hultman, Ruth J.F. Loos, Lili Milani, Aarno Palotie, Carlos Pato, Michele Pato, Danish Saleheen, Patrick F. Sullivan, Jessica Alföldi, Paul Cannon, Daniel G. MacArthur

**Affiliations:** National Heart & Lung Institute and MRC London Institute of Medical Sciences, Imperial College London, London, UK; Cardiovascular Research Centre, Royal Brompton & Harefield Hospitals NHS Trust, London, UK; Program in Medical and Population Genetics, Broad Institute of MIT and Harvard, Cambridge, MA 02142, USA; Analytic and Translational Genetics Unit, Massachusetts General Hospital, Boston, MA 02114, USA; 23andMe, Inc.; Program in Metabolism, Broad Institute of MIT and Harvard, Cambridge, MA 02142, USA; Center for Genomic Medicine, Massachusetts General Hospital, Boston, MA 02114, USA; Division of Endocrinology and Center for Basic and Translational Obesity Research, Boston Children’s Hospital, Boston, MA, USA; Program in Bioinformatics and Integrative Genomics, Harvard Medical School, Boston, MA 02115, USA; Program in Biological and Biomedical Sciences, Harvard Medical School, Boston, MA, 02115, USA; Michael J. Fox Foundation; Institute for Molecular Medicine Finland (FIMM), HiLIFE, University of Helsinki, Helsinki, Finland; National Institute for Health and Welfare, 00271, Helsinki, Finland; Department of Medical Epidemiology and Biostatistics, Karolinska Institutet, Stockholm, Sweden; Department of Biostatistics and Epidemiology, Perelman School of Medicine at the University of Pennsylvania, Philadelphia, PA, USA; The Charles Bronfman Institute for Personalized Medicine, Icahn School of Medicine at Mount Sinai, New York, NY, USA; Department of Medicine, Icahn School of Medicine at Mount Sinai, New York, NY, USA; Department of Psychiatry and the Behavioral Sciences, State University of New York, Downstate Medical Center, New York, NY, USA; Stanley Center for Psychiatric Research, Broad Institute of MIT and Harvard, Cambridge, Massachusetts 02142, USA; Estonian Genome Center, Institute of Genomics, University of Tartu, Tartu, Estonia; The Mindich Child Health and Development Institute, Icahn School of Medicine at Mount Sinai, New York, NY, USA; Department of Medicine, Perelman School of Medicine at the University of Pennsylvania, Philadelphia, PA, USA; Center for Non-Communicable Diseases, Karachi, Pakistan; Departments of Genetics and Psychiatry, University of North Carolina, Chapel Hill, NC, USA; Program in Medical and Population Genetics, Broad Institute of MIT and Harvard, Cambridge, Massachusetts 02142, USA; Analytic and Translational Genetics Unit, Massachusetts General Hospital, Boston, Massachusetts 02114, USA; European Molecular Biology Laboratory, European Bioinformatics Institute, Wellcome Genome Campus, Hinxton, Cambridge, CB10 1SD, United Kingdom; Data Sciences Platform, Broad Institute of MIT and Harvard, Cambridge, Massachusetts 02142, USA; Genomics Platform, Broad Institute of MIT and Harvard, Cambridge, Massachusetts 02142, USA; Broad Genomics, Broad Institute of MIT and Harvard, Cambridge, Massachusetts 02142, USA; Division of Genetics and Genomics, Boston Children’s Hospital, Boston, Massachusetts 02115, USA; Department of Pediatrics, Harvard Medical School, Boston, Massachusetts 02115, USA; Wellcome Sanger Institute, Wellcome Genome Campus, Hinxton, Cambridge CB10 1SA, UK; National Heart & Lung Institute and MRC London Institute of Medical Sciences, Imperial College London, London UK; Cardiovascular Research Centre, Royal Brompton & Harefield Hospitals NHS Trust, London UK; Unidad de Investigacion de Enfermedades Metabolicas. Instituto Nacional de Ciencias Medicas y Nutricion. Mexico City; Peninsula College of Medicine and Dentistry, Exeter, UK; Division of Preventive Medicine, Brigham and Women’s Hospital, Boston, Massachusetts, USA; Division of Cardiovascular Medicine, Brigham and Women’s Hospital and Harvard Medical School, Boston, Massachusetts, USA; Department of Cardiology, University Hospital, 43100 Parma, Italy; Department of Biology, Faculty of Natural Sciences, University of Haifa, Haifa, Israel; Departments of Medicine and Genetics, Albert Einstein College of Medicine, Bronx, NY, USA, 10461; Department of Quantitative Health Sciences, Lerner Research Institute, Cleveland Clinic, Cleveland, OH 44122, USA; Sorbonne Université, APHP, Gastroenterology Department, Saint Antoine Hospital, Paris, France; NHLBI and Boston University’s Framingham Heart Study, Framingham, Massachusetts, USA; Department of Medicine, Boston University School of Medicine, Boston, Massachusetts, USA; Department of Epidemiology, Boston University School of Public Health, Boston, Massachusetts, USA; Department of Biostatistics and Center for Statistical Genetics, University of Michigan, Ann Arbor, Michigan 48109; National Human Genome Research Institute, National Institutes of Health, Bethesda, MD, USA; The Charles Bronfman Institute for Personalized Medicine, Icahn School of Medicine at Mount Sinai, New York, NY; Department of Biochemistry, Wake Forest School of Medicine, Winston-Salem, NC, USA; Center for Genomics and Personalized Medicine Research, Wake Forest School of Medicine, Winston-Salem, NC, USA; Center for Diabetes Research, Wake Forest School of Medicine, Winston-Salem, NC, USA; Department of Cardiovascular Sciences, University of Leicester, Leicester, UK; NIHR Leicester Biomedical Research Centre, Glenfield Hospital, Leicester, UK; Department of Epidemiology and Biostatistics, Imperial College London, London, UK; Department of Cardiology, Ealing Hospital NHS Trust, Southall, UK; Imperial College Healthcare NHS Trust, Imperial College London, London, UK; Department of Medicine and Therapeutics, The Chinese University of Hong Kong, Hong Kong, China; Department of Medicine, Harvard Medical School, Boston, MA; Departments of Cardiovascular Medicine, Cellular and Molecular Medicine, Molecular Cardiology, and Quantitative Health Sciences, Cleveland Clinic, Cleveland, Ohio, USA; McLean Hospital, Belmont, MA; Department of Medicine, University of Mississippi Medical Center, Jackson, Mississippi, USA; Department of Epidemiology, Colorado School of Public Health, Aurora, Colorado, USA; Program in Medical and Population Genetics, Broad Institute of MIT and Harvard, Cambridge, MA, USA; Stanley Center for Psychiatric Research, Broad Institute of MIT and Harvard, Cambridge, MA, USA; Department of Medicine and Pharmacology, University of Illinois at Chicago; Department of Genetics, Texas Biomedical Research Institute, San Antonio, TX, USA; Department of Biostatistics, Boston University School of Public Health, Boston, MA 02118, USA; National Heart, Lung, and Blood Institute’s Framingham Heart Study, Framingham, MA 01702, USA; Cardiac Arrhythmia Service and Cardiovascular Research Center, Massachusetts General Hospital, Boston, MA; Cardiovascular Epidemiology and Genetics, Hospital del Mar Medical Research Institute (IMIM). Barcelona, Catalonia, Spain; CIBER CV, Barcelona, Catalonia, Spain; Departament of Medicine, Medical School, University of Vic-Central University of Catalonia. Vic, Catalonia, Spain; Institute for Cardiogenetics, University of Lübeck, Lübeck, Germany; 1. DZHK (German Research Centre for Cardiovascular Research), partner site Hamburg/Lübeck/Kiel, 23562 Lübeck, Germany; University Heart Center Lübeck, 23562 Lübeck, Germany; Helsinki University and Helsinki University Hospital, Clinic of Gastroenterology, Helsinki, Finland; Diabetes Unit and Center for Genomic Medicine, Massachusetts General Hospital; Programs in Metabolism and Medical & Population Genetics, Broad Institute; Department of Medicine, Harvard Medical School; Institute of Clinical Molecular Biology (IKMB), Christian-Albrechts-University of Kiel, Kiel, Germany; Bioinformatics Program, MGH Cancer Center and Department of Pathology; Cancer Genome Computational Analysis, Broad Institute; Endocrinology and Metabolism Department, Hadassah-Hebrew University Medical Center, Jerusalem, Israel; Department of Psychiatry and Behavioral Sciences; SUNY Upstate Medical University; Institute for Genomic Medicine, Columbia University Medical Center, Hammer Health Sciences, 1408, 701 West 168th Street, New York, New York 10032, USA; Department of Genetics & Development, Columbia University Medical Center, Hammer Health Sciences, 1602, 701 West 168th Street, New York, New York 10032, USA; Centro de Investigacion en Salud Poblacional. Instituto Nacional de Salud Publica MEXICO; Lund University, Sweden; Lund University Diabetes Centre; Human Genetics Center, University of Texas Health Science Center at Houston, Houston, TX 77030; Department of Neurology, Columbia University; Institute of Genomic Medicine, Columbia University; Institute of Biomedicine, University of Eastern Finland, Kuopio, Finland; Department of Psychiatry, PL 320, Helsinki University Central Hospital, Lapinlahdentie, 00 180 Helsinki, Finland; Icahn School of Medicine at Mount Sinai, New York, NY, USA; Department of Neurology, Helsinki University Central Hospital, Helsinki, Finland; Department of Public Health, Faculty of Medicine, University of Helsinki, Finland; Center for Genomic Medicine, Massachusetts General Hospital, Boston, Massachusetts 02114, USA; Cardiovascular Disease Initiative and Program in Medical and Population Genetics, Broad Institute of MIT and Harvard, Cambridge, Massachusetts 02142, USA; Center for Genome Science, Korea National Institute of Health, Chungcheongbuk-do, Republic of Korea; MRC Centre for Neuropsychiatric Genetics & Genomics, Cardiff University School of Medicine, Hadyn Ellis Building, Maindy Road, Cardiff CF24 4HQ; National Heart and Lung Institute, Cardiovascular Sciences, Hammersmith Campus, Imperial College London, London, UK; Department of Health, THL-National Institute for Health and Welfare, 00271 Helsinki, Finland; Section of Cardiovascular Medicine, Department of Internal Medicine, Yale School of Medicine, New Haven, Connecticut3Center for Outcomes Research and Evaluation, Yale-New Haven Hospital, New Haven, Connecticut; Division of Pediatric Gastroenterology, Emory University School of Medicine, Atlanta, Georgia, USA; Department of Internal Medicine, Seoul National University Hospital, Seoul, Republic of Korea; The University of Eastern Finland, Institute of Clinical Medicine, Kuopio, Finland; Kuopio University Hospital, Kuopio, Finland; Department of Clinical Chemistry, Fimlab Laboratories and Finnish Cardiovascular Research Center-Tampere, Faculty of Medicine and Health Technology, Tampere University, Finland; The Mindich Child Health and Development Institute, Icahn School of Medicine at Mount Sinai, New York, NY; Li Ka Shing Institute of Health Sciences, The Chinese University of Hong Kong, Hong Kong, China; Hong Kong Institute of Diabetes and Obesity, The Chinese University of Hong Kong, Hong Kong, China; Cardiovascular Research REGICOR Group, Hospital del Mar Medical Research Institute (IMIM). Barcelona, Catalonia; Department of Genetics, Harvard Medical School, Boston, MA, USA; Oxford Centre for Diabetes, Endocrinology and Metabolism, University of Oxford, Churchill Hospital, Old Road, Headington, Oxford, OX3 7LJ UK; Wellcome Centre for Human Genetics, University of Oxford, Roosevelt Drive, Oxford OX3 7BN, UK; Oxford NIHR Biomedical Research Centre, Oxford University Hospitals NHS Foundation Trust, John Radcliffe Hospital, Oxford OX3 9DU, UK; F Widjaja Foundation Inflammatory Bowel and Immunobiology Research Institute, Cedars-Sinai Medical Center, Los Angeles, CA, USA; Atherogenomics Laboratory, University of Ottawa Heart Institute, Ottawa, Canada; Division of General Internal Medicine, Massachusetts General Hospital, Boston, MA, 02114; Program in Population and Medical Genetics, Broad Institute, Cambridge, MA; Department of Clinical Sciences, University Hospital Malmo Clinical Research Center, Lund University, Malmo, Sweden; Lund University, Dept. Clinical Sciences, Skane University Hospital, Malmo, Sweden; Instituto Nacional de Medicina Genómica (INMEGEN), Mexico City, 14610, Mexico; Medical Research Institute, Ninewells Hospital and Medical School, University of Dundee, Dundee, UK; Department of Molecular Medicine and Biopharmaceutical Sciences, Graduate School of Convergence Science and Technology, Seoul National University, Seoul, Republic of Korea; Department of Psychiatry, Keck School of Medicine at the University of Southern California, Los Angeles, California, USA; Department of Psychiatry and Behavioral Sciences, Johns Hopkins University School of Medicine, Baltimore, Maryland, USA; Division of Genetics and Epidemiology, Institute of Cancer Research, London SM2 5NG; Medical Research Center, Oulu University Hospital, Oulu, Finland and Research Unit of Clinical Neuroscience, Neurology, University of Oulu, Oulu, Finland; Research Center, Montreal Heart Institute, Montreal, Quebec, Canada, H1T 1C8; Department of Medicine, Faculty of Medicine, Université de Montréal, Québec, Canada; Broad Institute of MIT and Harvard, Cambridge MA, USA; Department of Biomedical Informatics, Vanderbilt University Medical Center, Nashville, Tennessee, USA; Department of Medicine, Vanderbilt University Medical Center, Nashville, Tennessee, USA; National Institute for Health and Welfare, Helsinki, Finland; Deutsches Herzzentrum München, Germany; Technische Universität München; Division of Cardiovascular Medicine, Nashville VA Medical Center and Vanderbilt University, School of Medicine, Nashville, TN 37232-8802, USA; Department of Psychiatry, Icahn School of Medicine at Mount Sinai, New York, NY, USA; Department of Genetics and Genomic Sciences, Icahn School of Medicine at Mount Sinai, New York, NY, USA; Institute for Genomics and Multiscale Biology, Icahn School of Medicine at Mount Sinai, New York, NY, USA; Institute of Clinical Medicine, neurology, University of Eastern Finad, Kuopio, Finland; Department of Twin Research and Genetic Epidemiology, King’s College London, London UK; Saw Swee Hock School of Public Health, National University of Singapore, National University Health System, Singapore; Department of Medicine, Yong Loo Lin School of Medicine, National University of Singapore, Singapore; Duke-NUS Graduate Medical School, Singapore; Life Sciences Institute, National University of Singapore, Singapore; Department of Statistics and Applied Probability, National University of Singapore, Singapore; Folkhälsan Institute of Genetics, Folkhälsan Research Center, Helsinki, Finland; HUCH Abdominal Center, Helsinki University Hospital, Helsinki, Finland; Center for Behavioral Genomics, Department of Psychiatry, University of California, San Diego; Institute of Genomic Medicine, University of California, San Diego; Juliet Keidan Institute of Pediatric Gastroenterology, Shaare Zedek Medical Center, The Hebrew University of Jerusalem, Israel; Instituto de Investigaciones Biomédicas UNAM Mexico City; Instituto Nacional de Ciencias Médicas y Nutrición Salvador Zubirán Mexico City; National Heart & Lung Institute & MRC London Institute of Medical Sciences, Imperial College London, London UK; Radcliffe Department of Medicine, University of Oxford, Oxford UK; Department of Gastroenterology and Hepatology, University of Groningen and University Medical Center Groningen, Groningen, the Netherlands; Department of Physiology and Biophysics, University of Mississippi Medical Center, Jackson, MS 39216, USA; Program in Infectious Disease and Microbiome, Broad Institute of MIT and Harvard, Cambridge, MA, USA; Center for Computational and Integrative Biology, Massachusetts General Hospital

## Abstract

Human genetic variants predicted to cause loss-of-function of protein-coding genes (pLoF variants) provide natural *in vivo* models of human gene inactivation, and can be valuable indicators of gene function and the potential toxicity of therapeutic inhibitors targeting these genes^1,2^. Gain-of-kinase-function variants in *LRRK2* are known to significantly increase the risk of Parkinson’s disease^3,4^, suggesting that inhibition of LRRK2 kinase activity is a promising therapeutic strategy. While preclinical studies in model organisms have raised some on-target toxicity concerns^5–8^, the biological consequences of LRRK2 inhibition have not been well-characterized in humans. Here we systematically analyse pLoF variants in *LRRK2* observed across 141,456 individuals sequenced in the Genome Aggregation Database (gnomAD)^9^, 49,960 exome sequenced individuals from the UK Biobank, and over 4 million participants in the 23andMe genotyped dataset. After stringent variant curation, we identify 1,455 individuals with high-confidence pLoF variants in *LRRK2*, 82.5% with experimental validation. We show that heterozygous pLoF variants in *LRRK2* reduce LRRK2 protein levels but are not strongly associated with reduced life expectancy, or with any specific phenotype or disease state. These data suggest that therapeutics that partially downregulate LRRK2 levels or kinase activity are unlikely to have major on-target safety liabilities. Our results demonstrate the value of large-scale genomic databases and phenotyping of human LoF carriers for target validation in drug discovery.

## Main text

Novel therapeutic strategies are desperately needed in Parkinson’s disease (PD), one of the most common age-related neurological diseases, which affects about 1% of people over the age of 60^10,11^. Around 30% of familial and 3-5% of sporadic PD cases have been linked to a genetic cause^12^. *LRRK2* missense variants account for a significant fraction of cases, including high-penetrance variants^13^, moderately penetrant variants such as G2019S^14^, and risk factors identified in genome-wide association studies^15^. Although the precise mechanism by which *LRRK2* variants mediate their pathogenicity remains unclear, a common feature is augmentation of kinase activity associated with disease-relevant alterations in cell models^3,16,17^. Discovery of Rab GTPases as substrates of LRRK2^18^ has also highlighted the role of LRRK2 in regulation of the endolysosomal and vesicular trafficking pathways implicated in Parkinson’s disease^18,19^. LRRK2 kinase activity is also upregulated more generally in PD patients (with and without LRRK2 variants)^20^. LRRK2 has therefore become a prominent drug target, with multiple LRRK2 kinase inhibitors and suppressors^21^ in development as disease-modifying treatments for PD^20,22,23^.

Despite these promising indications, there are concerns about the potential toxicity of LRRK2 inhibitors. These mainly arise from preclinical studies, where homozygous knockouts of *LRRK2* in mice, and high dose toxicology studies of LRRK2 kinase inhibitors in rats and primates, have shown abnormal phenotypes in lung, kidney and liver^5–8^. While model organisms have been invaluable in understanding the function of LRRK2, they also have important limitations as exemplified by the inconsistencies in the phenotypic consequences of reduced LRRK2 activity seen among yeast, fruit flies, worms, mice, rats and non-human primates^24^. Complementary data from natural human knockouts are critical for understanding both gene function and the potential consequences of a long-term reduction in LRRK2 kinase activity in humans.

Large-scale human genetics is an increasingly powerful source of data for the discovery and validation of therapeutic targets in humans^1^. Predicted loss-of-function (pLoF) variants, predicted to largely or entirely abolish the function of affected alleles, are a particularly informative class of genetic variation. Such variants are natural models for life-long organism-wide inhibition of the target gene, and can provide information about both the efficacy and safety of a candidate target^2,25–28^. However, pLoF variants tend to be rare in human populations^29^, and are also enriched for both sequencing and annotation artefacts^30^. As such, leveraging pLoF variation in drug target assessment typically requires very large collections of genetically and phenotypically characterized individuals, combined with deep curation of the target gene and candidate variants^31^. Although previous studies of pLoF variants in *LRRK2* have found no association with PD risk^32^, no study has assessed the broader phenotypic consequences of such variants.

We identified *LRRK2* pLoF variants, and assessed the associated phenotypic changes, in three large cohorts of genetically characterized individuals. Firstly, we annotated *LRRK2* pLoF variants in two large sequencing cohorts: the gnomAD v2.1.1 dataset, which contains 125,748 exomes and 15,708 genomes from unrelated individuals^9^, and the 46,062 exome-sequenced unrelated European individuals from the UK Biobank^33^. We identified 633 individuals in gnomAD and 258 individuals in the UK Biobank with 150 unique candidate *LRRK2* LoF variants, a combined carrier frequency of 0.48%. All variants were observed only in the heterozygous state. Compared to the spectrum observed across all genes, *LRRK2* is not significantly depleted for LoF variants in gnomAD (loss-of-function observed/expected upper bound fraction (LOEUF)^9^=0.64).

To ensure that the 150 identified variants are indeed expected to cause LoF, we manually curated them to remove those of low-quality or with annotation errors (Figure 1a; Supplementary Tables 1 and 2). We removed 16 variants identified either as low-confidence by the Loss-of-function Transcript Effect Estimator (LOFTEE; 6 variants in 409 individuals)^9^, or manually curated as low-quality or unlikely to cause true LoF (10 variants in 129 individuals; see methods). One additional individual was excluded from the UK Biobank cohort as they carried both an pLoF variant and the G2019S risk allele.

**Figure 1:**
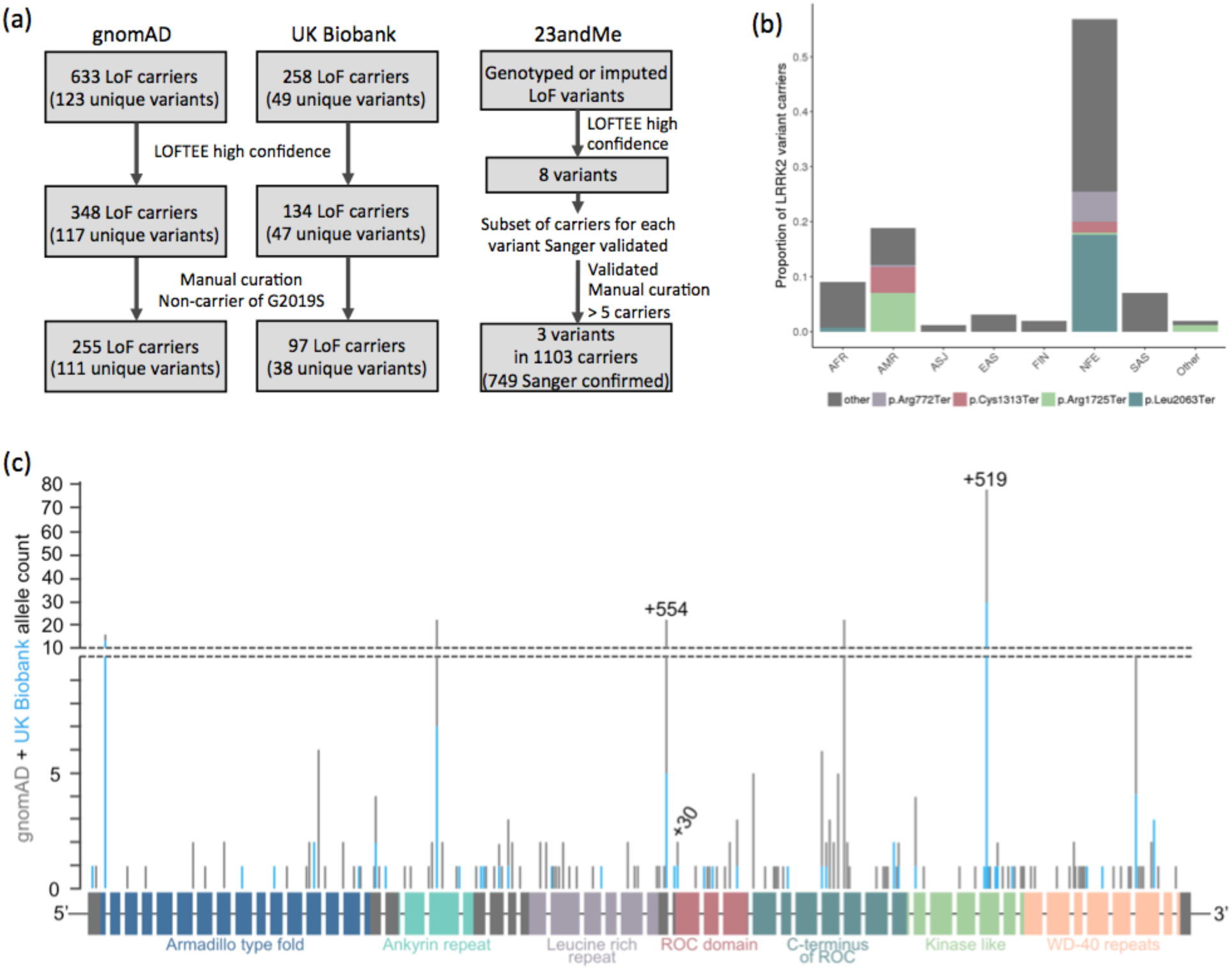
Annotation and curation of candidate *LRRK2* pLoF variants. a) Flow chart showing the variant filtering and curation of candidate *LRRK2* LoF variants in the gnomAD, UK Biobank and 23andMe cohorts. Of the 1,103 carriers identified in 23andMe, 749 were confirmed by Sanger sequencing with the remainder untested. b) The ancestry distribution of *LRRK2* pLoF variant carriers in gnomAD (AFR - African/African American; AMR - American/Latino; ASJ - Ashkenazi Jewish; EAS - East Asian; FIN - Finnish; NFE - Non-Finnish European; SAS - South Asian). pLoF variants seen more than 10 times appear in colour with remaining variants in grey. *LRRK2* pLoF variants are mostly individually extremely rare (less than 1/10,000 carrier frequency), with the exception of two nonsense variants almost exclusively restricted to the admixed American/Latino population (Cys1313Ter and Arg1725Ter), and two largely Non-Finnish European-specific variants (Leu2063Ter and Arg772Ter). All variant protein descriptions are with respect to ENSP00000298910.7. c) Schematic of the LRRK2 gene with pLoF variants marked by position, with the height of the marker corresponding to allele count in gnomAD (grey bars) and UK Biobank (blue bars). The 51 exons are shown as rectangles coloured by protein domain, with the remaining exons in grey. The three variants genotyped in the 23andMe cohort are annotated with their sample count in black text.

This analysis results in a final dataset of 255 gnomAD individuals and 97 UK biobank individuals with 134 unique high-confidence pLoF variants (Figure 1a), an overall carrier frequency of 0.19%; less than half the frequency estimated from uncurated variants, reaffirming the importance of thorough curation of candidate LoF variants^31^. A subset of 25 gnomAD samples with 19 unique *LRRK2* pLoF variants with DNA available were all successfully validated by Sanger sequencing (Supplementary Table 3).

Secondly, we examined *LRRK2* pLoF variants in over four million consented research participants from the personal genetics company 23andMe, Inc. Participants have been genotyped on a variety of platforms, and imputed against a reference panel comprising 56.5 million variants from 1000 Genomes phase 3^34^, and UK10K^35^. Eight putative *LRRK2* LoF variants were identified in the 23andMe cohort (LOFTEE high-confidence). After manual curation all putative carriers of each variant were submitted for validation by Sanger sequencing. Variants with <5 confirmed carriers were excluded from the analysis. The resulting cohort consisted of 749 individuals, each of whom is a Sanger-confirmed carrier for one of three pLoF variants (Figure 1a; Supplementary Table 4). The high rate of Sanger confirmation for rs183902574 (>98%) allowed confident addition of 354 putative carriers of rs183902574, from expansion of the 23andMe dataset, without Sanger confirmation. Analyses with and without these genotyped only carriers were not significantly different (Supplementary Table 6). Across the two most frequent pLoF alleles we observed an extremely small number (<5) of sequence-confirmed homozygotes; however, given the very small number of observations, we can make no robust inference except that homozygous inactivation of LRRK2 appears to be compatible with life. For the remainder of this manuscript we focus on heterozygous pLoF carriers.

The three combined datasets provide a total of 1,455 carrier individuals with 134 unique *LRRK2* pLoF variants. These variants are found across all major continental populations (Figure 1b; Supplementary Figure 1) and show neither any obvious clustering along the length of the LRRK2 protein, nor specific enrichment or depletion in any of the known annotated protein domains (Chi-square *P*=0.22; Figure 1c), consistent with signatures of true LoF^31^.

To confirm that *LRRK2* pLoF variants lead to a reduction in LRRK2 protein levels, we analysed total protein lysates from cell lines with three unique pLoF variants by immunoblotting. We obtained lymphoblastoid cell lines (LCLs) from two families with naturally occuring heterozygous LoF variants, and a third variant was CRISPR/Cas9 engineered into embryonic stem cells (Supplementary Figure 2), which were differentiated into cardiomyocytes. In all instances, LRRK2 protein levels were visibly reduced compared to non-carrier controls (Figure 2). These results agree with a previous study which assessed three separate pLoF variants, and found significant reductions in LRRK2 protein levels^32^. Together, these six functionally validated variants represent 82.5% of total pLoF carriers in this study (1,201/1,455). Although heterozygous pLoF carriers have LRRK2 protein remaining, we believe that this state represents the closest genetic model to the partial kinase inactivation expected to be generated by therapeutic LRRK2 inhibitors.

**Figure 2:**
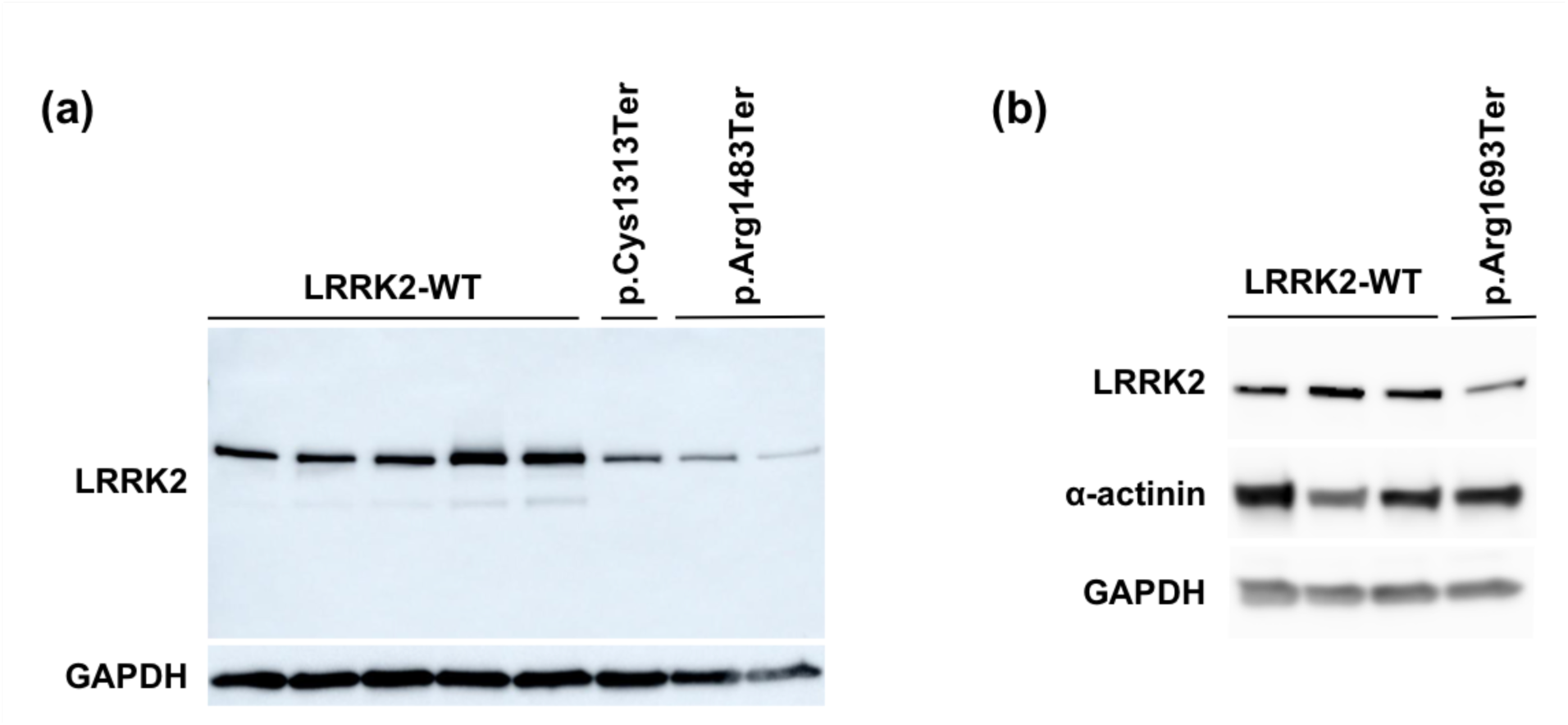
*LRRK2* pLoF heterozygotes have reduced LRRK2 protein compared to cells harboring no LoF variants. a) Immunoblot of LRRK2 and loading control GAPDH on lymphoblastoid cell lines from five individuals harboring no pLoF variants (LRRK2-WT) and three individuals harboring a heterozygous pLoF variant (Cys1313Ter - 12-40699748-T-A; Arg1483Ter - 12-40704362-C-T). b) Immunoblot of LRRK2, α-actinin (specific to muscle), and GAPDH on three control lines and one CRISPR/Cas9 engineered LRRK2 heterozygous line of cardiomyocytes differentiated from ESCs (Arg1693Ter - 12-40714897-C-T). All variant protein descriptions are with respect to ENSP00000298910.7.

We next sought to determine whether lifelong lowering of LRRK2 protein levels through LoF results in an apparent reduction in lifespan. We found no significant difference between the age distribution of *LRRK2* pLoF variant carriers and non-carriers in either the gnomAD or 23andMe datasets (two-sided Kolmogorov-Smirnov *P*=0.085 and 0.46 respectively; Figure 3a), suggesting no major impact on longevity, though we note that at current sample sizes we are only powered to detect a strong effect (see Methods).

**Figure 3:**
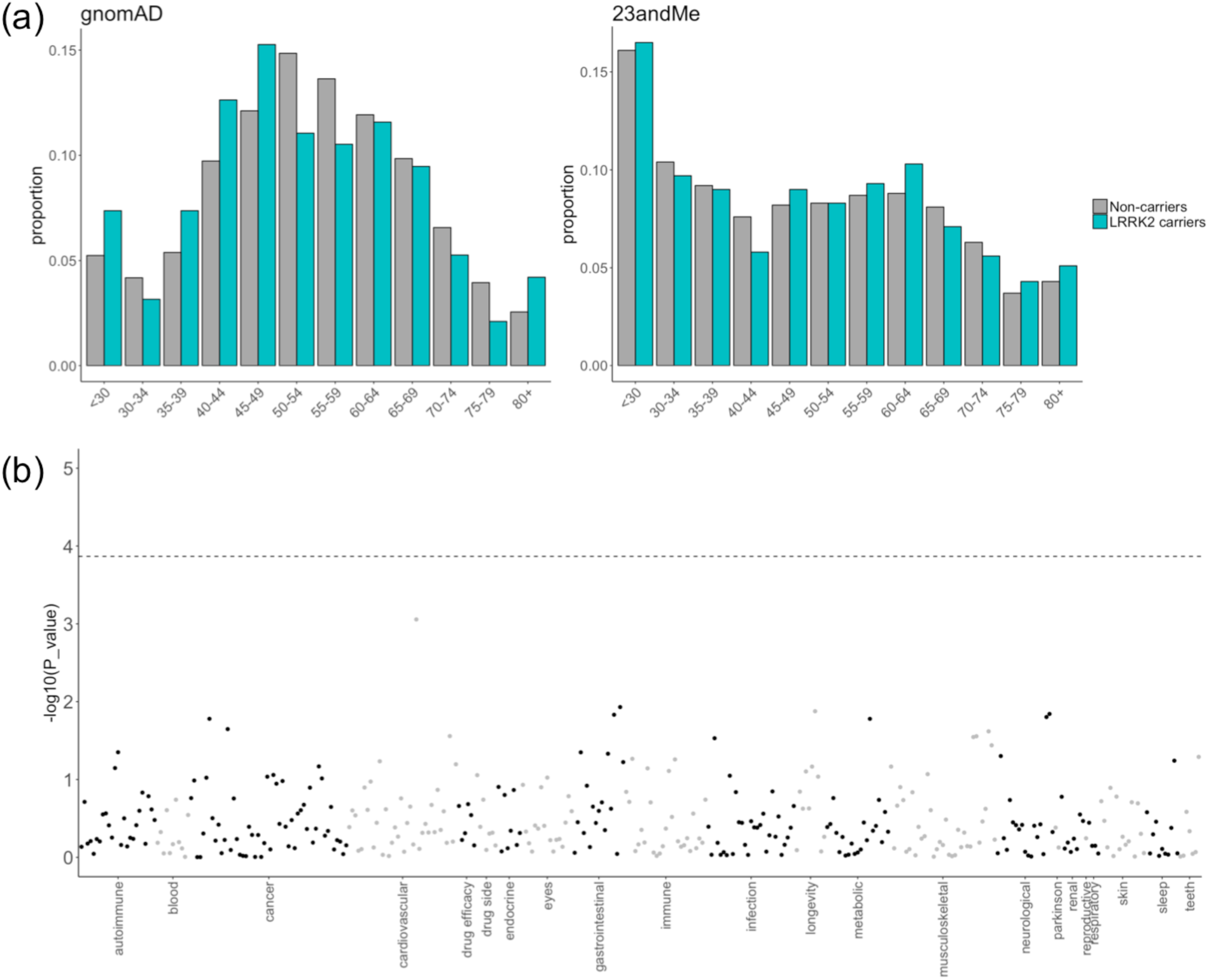
*LRRK2* pLoF variants are not associated with reduced lifespan and are not associated with any adverse phenotypes. (a) The age distributions of *LRRK2* pLoF carriers are not significantly different from those of non-carriers in both gnomAD and 23andMe. (b) Manhattan plot of phenome-wide association study results for carriers of three *LRRK2* pLoF variants against non-carriers in the 23andMe cohort. Each point represents a distinct phenotype with these grouped into related categories (delineated by alternating black and grey points). The dotted horizontal line represents a Bonferoni corrected *P* value threshold for 368 tests.

For a subset of studies within gnomAD, phenotype data is available from study or national biobank questionnaires or from linked electronic health records (see online methods). We manually reviewed these records for all 60 of the 255 gnomAD *LRRK2* pLoF carriers for whom data were available, and recorded any phenotypes affecting the lung, liver, kidney, cardiovascular system, nervous system, immunity, and cancer (Supplementary Table 5). We did not find an over-representation of any particular phenotype or phenotype category in *LRRK2* pLoF carriers.

The 23andMe dataset includes self-reported data for thousands of phenotypes across a diverse range of categories. We performed a phenome-wide association study on *LRRK2* pLoF carriers compared to non-carriers for 368 health-related traits (see methods) and found no significant association between any individual phenotype and carrier status (Figure 3b). In particular, we found no significant associations with any lung, liver or kidney phenotypes (Supplementary Tables 6 and 7).

The UK Biobank resource includes measurements for 30 blood serum and four urine biomarkers. We found no difference in any of these biomarkers between pLoF carriers and controls (Supplementary Table 8; Supplementary Figure 3). In particular, there was no difference between carriers and non-carriers for urine biomarkers transformed into clinical measures of kidney function (Figure 4a; see online methods) and no difference in six blood biomarkers commonly used to assess liver function (Figure 4c). We also observed no difference in spirometry measurements of lung function (Figure 4b).

**Figure 4:**
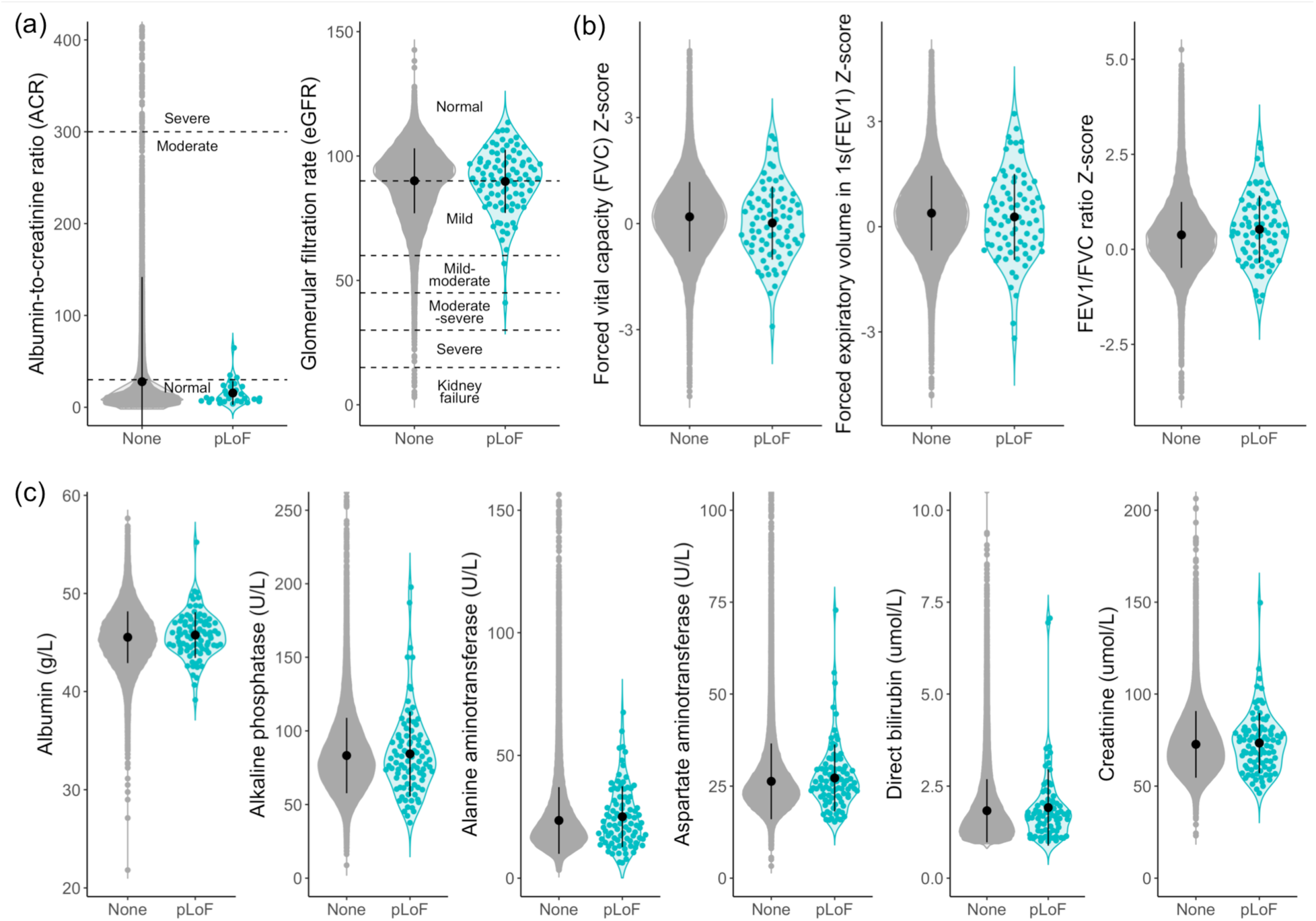
*LRRK2* pLoF carriers do not have impaired lung, liver or kidney function. For all plots, pLoF carriers are shown in teal and non-carriers in grey. The mean and 1 standard deviation are represented by the black circle and line. (a) Urine biomarkers Albumin and Creatinine transformed into two clinical markers of kidney function (see online methods). No pLoF carriers show signs of severely impaired kidney function. (b) Z-scores of age, sex and height corrected spirometry measures of lung function^36^. (c) Blood serum biomarkers of liver function. The plots for alkaline phosphatase, alanine aminotransferase, aspartate aminotransferase, bilirubin and creatinine have been top truncated removing 47, 29, 92, 8 and 27 non-carriers respectively. The violin plots and summary statistics were calculated on the full dataset. All pLoF carriers are within each plot area.

We grouped self-reported disease diagnoses in UK Biobank individuals into categories corresponding to organ system and/or mechanism (Supplementary Table 9; see online methods). We observed no enrichment for any of these phenotype groups in *LRRK2* pLoF carriers when compared to non-carriers (Supplementary Table 10). We also mined ICD10 codes from hospital admissions and death records for any episodes relating to lung, liver and kidney phenotypes, removing any with a likely infectious or other external causes (Supplementary Table 11; see online methods), and identified six pLoF carriers with ICD10 codes relating to these organ systems (6.19%), compared to 4,536 non-carriers (9.87%; Supplementary Tables 12 and 13).

Our results indicate that approximately one in every 500 humans is heterozygous for a pLoF variant in *LRRK2*, resulting in a systemic lifelong decrease in LRRK2 protein levels, and that this partial inhibition has no discernible effect on survival or health. These results suggest that drugs that reduce LRRK2 kinase activity are unlikely to show toxicity due to on-target effects. This observation is particularly important given that individuals at risk of PD will need to take therapeutics chronically, likely soon after clinical diagnosis and extending for decades. While initial phase I studies of the safety of LRRK2 kinase inhibitors have shown a promising safety profile^23,37^, these are not yet able to address the question of long-term, on-target pharmacology-related safety profiles.

The rarity of pLoF variants in *LRRK2*, combined with the relatively low prevalence of PD, prevents us from directly assessing whether LRRK2 inhibition may reduce the incidence of PD with current sample sizes (Supplementary Table 6). Future cohorts with much larger numbers of sequenced and phenotyped individuals (likely in the millions of samples) will be required to answer this question. As such, our study focuses entirely on the question of whether partial genetic LRRK2 inactivation has broader phenotypic consequences that might correspond to adverse effects of chronic inhibition of LRRK2 inhibitors.

We acknowledge multiple limitations to the work described here. Firstly, we relied on heterogeneous phenotype data mostly derived from self-reported questionnaires. Both 23andMe and gnomAD record only age at recruitment, and participants are of a relatively young age compared to the typical age of onset for PD. Our ascertainment of *LRRK2* pLoF variants in 23andMe was necessarily incomplete, due to the availability of targeted genotyping rather than sequencing data in this cohort; this means that a subset of the 23andMe individuals treated as non-carriers could be carriers of *LRRK2* pLoF variants not genotyped or imputed in this dataset. Finally, the low-frequency of naturally occuring pLoF variants in *LRRK2* results in a relatively small number of carriers which could be assessed for each biomarker and phenotype, meaning that we are not well-powered to detect subtle or infrequent clinical phenotypes arising from LRRK2 haploinsufficiency. However, our study strongly suggests that any clinical phenotype arising from partial inactivation of LRRK2 is likely to be substantially more benign than early-onset PD.

This study provides an important proof of principle for the value of very large genetically and phenotypically characterized cohorts, combined with thorough variant curation, in exploring the safety profile of candidate drug targets. Over the coming years, the availability of complete exome or genome sequence data for hundreds of thousands of individuals who are deeply phenotyped and/or available for genotype-based recontact studies, combined with deep curation and experimental validation of candidate pLoF variants, will provide powerful resources for therapeutic target validation as well as broader studies of the biology of human genes.

## Data and code availability

The data presented in this manuscript and the code used to make the figures are available in https://github.com/macarthur-lab/gnomad_lrrk2.

## Acknowledgments

We thank the research participants and employees of 23andMe, Inc., and the research participants in gnomAD, for making this work possible. N.W. is supported by a Rosetrees and Stoneygate Imperial College Research Fellowship. E.V.M. is supported by NIH F31 AI22592. KJK was supported by NIGMS F32 GM115208. This work was supported by the Michael J. Fox Foundation for Parkinson’s Research grant 12868, NIDDK U54DK105566, NIGMS R01GM104371, Wellcome Trust [107469/Z/15/Z]; Medical Research Council (UK); NIHR Royal Brompton Biomedical Research Unit; NIHR Imperial Biomedical Research Centre. T.E is supported by Estonian Research Council grant PUT1660. L.M. is supported by Estonian Research Council grant PRG184. This research has been conducted using the UK Biobank Resource (https://doi.org/10.1101/572347) under Application Number 42890.

## Conflicts of interest

A.K., B.A., A.G., P.M., P.C., and members of the 23andMe Research Team are current or former employees of 23andMe, Inc., and hold stock or stock options in 23andMe. DGM is a founder with equity in Goldfinch Bio, and has received research support from AbbVie, Astellas, Biogen, BioMarin, Eisai, Merck, Pfizer, and Sanofi-Genzyme. EVM has received research support in the form of charitable contributions from Charles River Laboratories and Ionis Pharmaceuticals, and has consulted for Deerfield Management. KJK owns stock in Personalis. MJD is a founder of Maze Therapeutics.

## Supplementary Tables and Figures

Supplementary Table 1: Curated *LRRK2* LoF variants identified in gnomAD. All positions are with respect to GRCh37.

Supplementary Table 2: Curated *LRRK2* LoF variants identified in 46,062 unrelated European individuals in the UK Biobank. All positions are shown with respect to both GRCh37 and GRCh38.

Supplementary Table 3: Details of variants and primers for Sanger validation of *LRRK2* LoF carriers. All positions are with respect to GRCh37.

Supplementary Table 4. Details of *LRRK2* variants identified in the 23andMe cohort. All positions are with respect to GRCh37. All variant protein descriptions are annotated on ENSP00000298910.

Supplementary Table 5. Summary of phenotype data from gnomAD *LRRK2* LoF carriers. All positions are with respect to GRCh37.

Supplementary Table 6: Association statistics for phenotypes specific to lung, kidney and liver (also plotted in Figure 3c). For privacy reasons, counts of less than five are rounded. None of the phenotypes associated with carrier status, even without correction (p < 0.05).

Supplementary Table 7: 23andMe phenotype association statistics for all phenotypes plotted in Figure 3b.

Supplementary Table 8: Details of 30 blood serum and 4 urine biomarkers tested in the UK Biobank dataset. Shown are the count of pLoF carriers and non-carriers as well as the mean and standard-deviation within each cohort. *P*-values correspond to a 2-side Wilcoxon test.

Supplementary Table 9: Details of phenotype groupings for diseases coded within the field ‘2002 Non-cancer illness code, self-reported’. Shown are the tree positions within UK Biobank along with primary and secondary groupings used in this study.

Supplementary Table 10: Association statistics for all phenotype groupings in the UK Biobank. pLoF carriers and non-carriers were counted only once corresponding to whether or not they had any of the self-reported phenotypes within a grouping. P-values correspond to a Fisher’s exact test.

Supplementary Table 11: ICD10 codes taken from UK Biobank fields ‘41270 - Diagnoses - ICD10’, ‘40001 - Underlying (primary) cause of death: ICD10’ and ‘40002 - Contributory (secondary) causes of death: ICD10’ to detect lung, liver and kidney disease episodes.

Supplementary Table 12: ICD10 codes relating to lung, liver and kidney disease reported in LRRK2 pLoF carriers.

Supplementary Table 13: Details of ICD10 codes relating to lung, liver and kidney disease reported in each of the six LRRK2 pLoF carriers with at least one reported. All positions are with respect to GRCh37.

Supplementary Table 14: Details of 30 blood serum and 4 urine biomarkers tested in the UK Biobank dataset. Shown are the count of pLoF carriers and G2019S risk allele carriers as well as the mean and standard-deviation within each cohort. P-values correspond to a 2-side Wilcoxon test.

Supplementary Table 15: The power to detect an age effect of the same magnitude as APOE rs429358 in the 23andMe dataset.

**Supplementary Figure 1.**
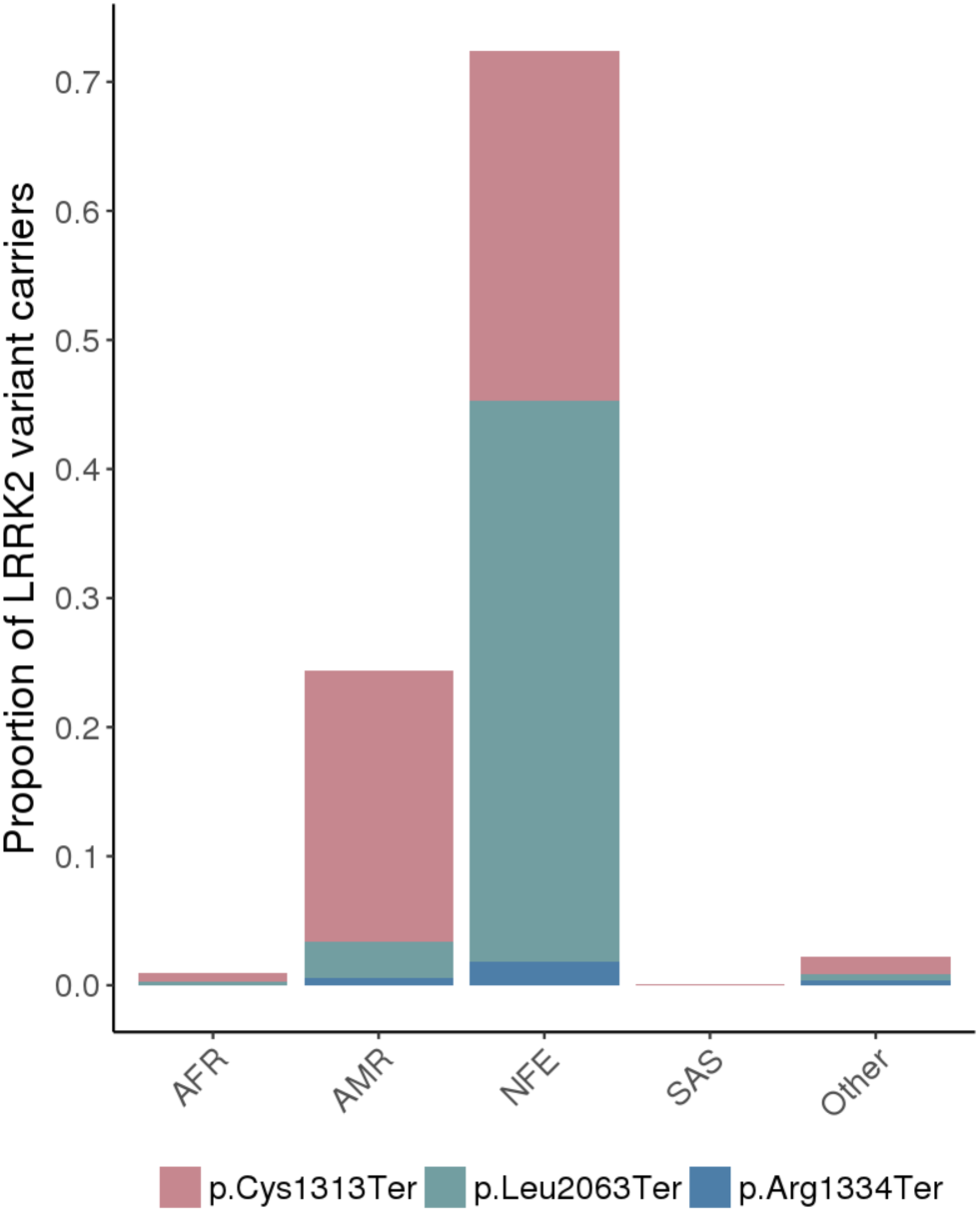
Ethnicity distribution of LRRK2 LoF carriers in the 23andMe cohort.

**Supplementary Figure 2.**
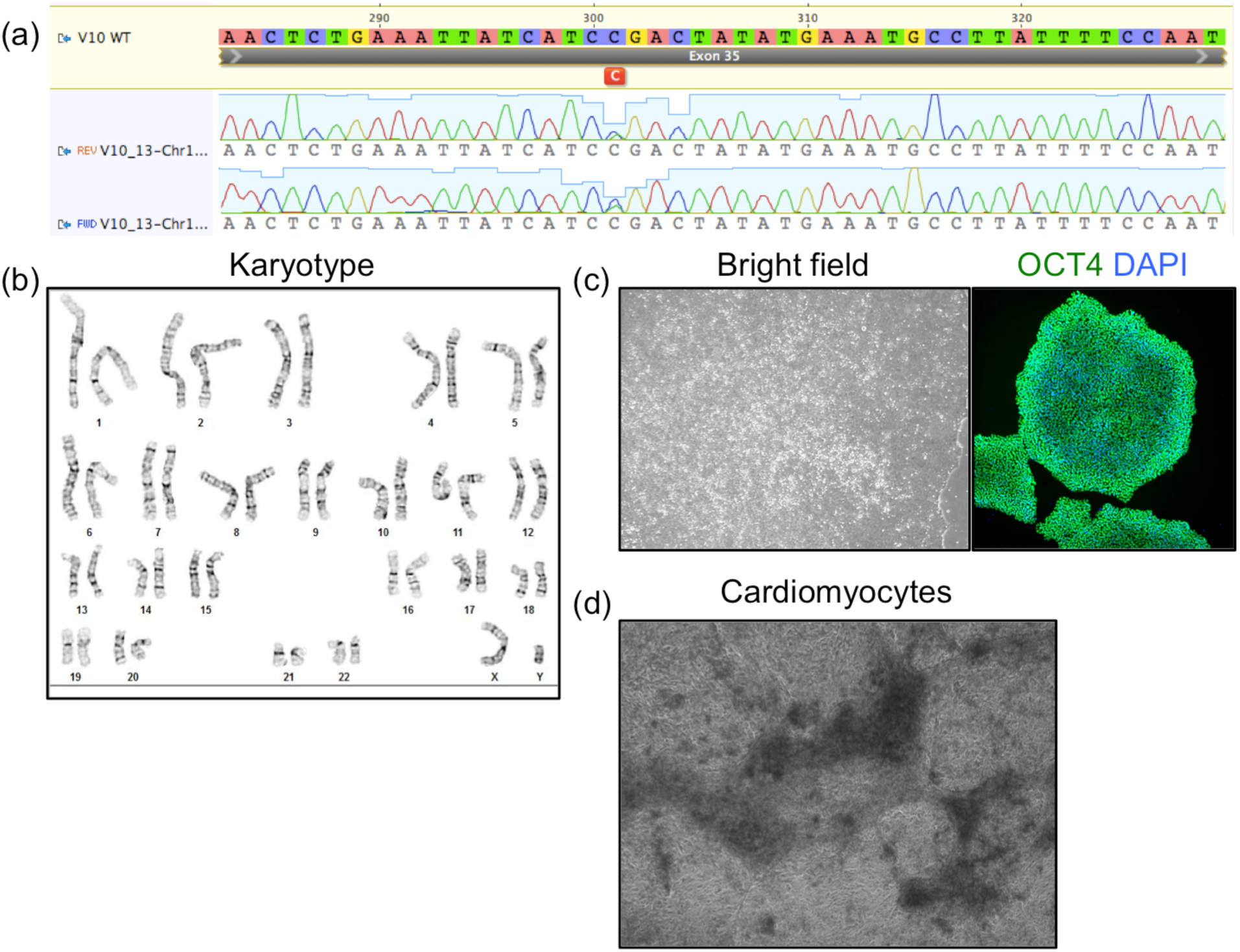
a) Sanger sequence of isogenic hESC engineered colony for heterozygous LRRK2 variant 10, clone 13 (GRCh37:12-40714897-C-T). The engineered cell line maintains b) a normal karyotype, c) normal colony morphology and expression of OCT4, and d) differentiates into the cardiomyocyte lineage. The bright field image of cardiomyocytes was captured at day 17 of the differentiation protocol.

**Supplementary Figure 3:**
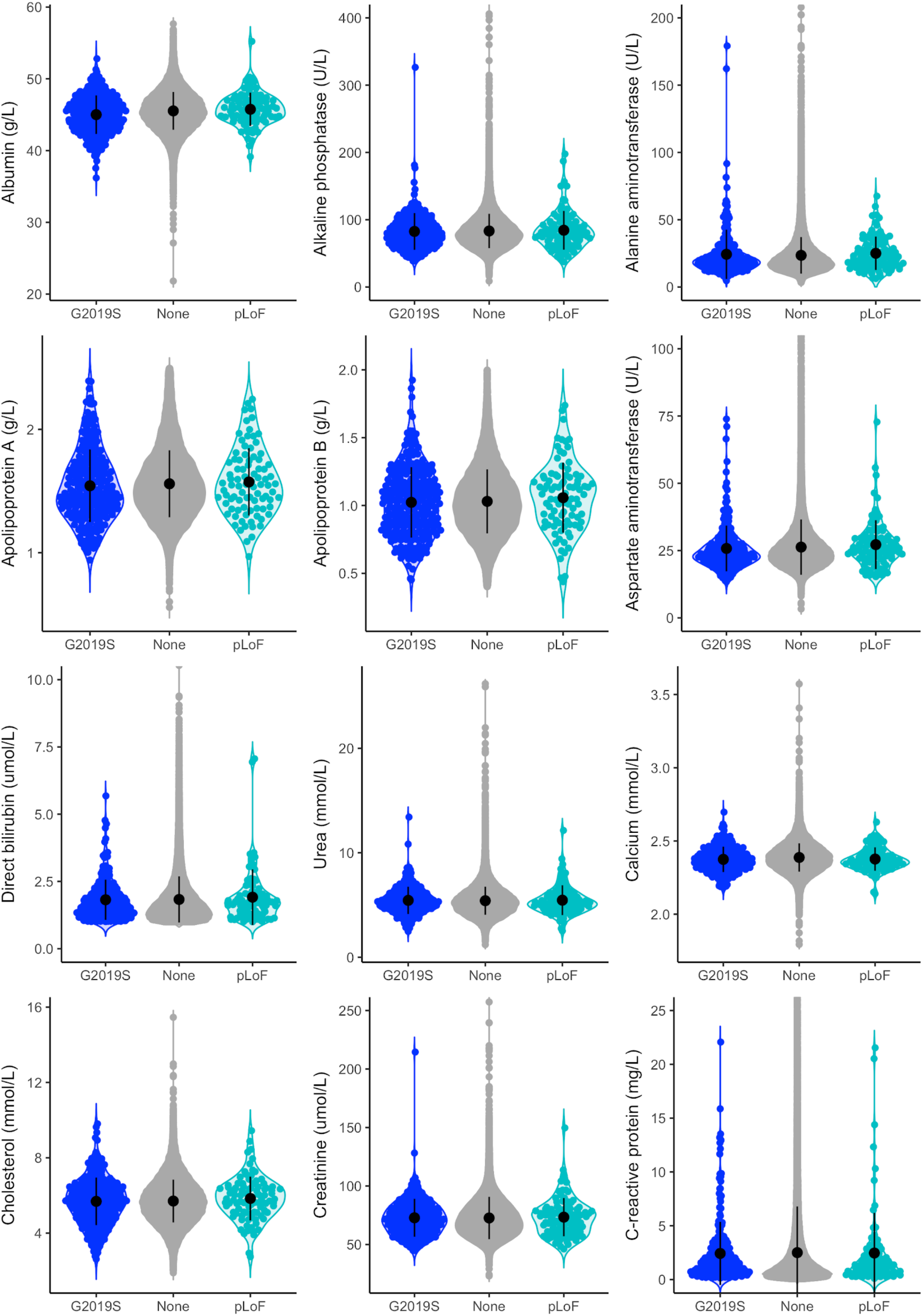
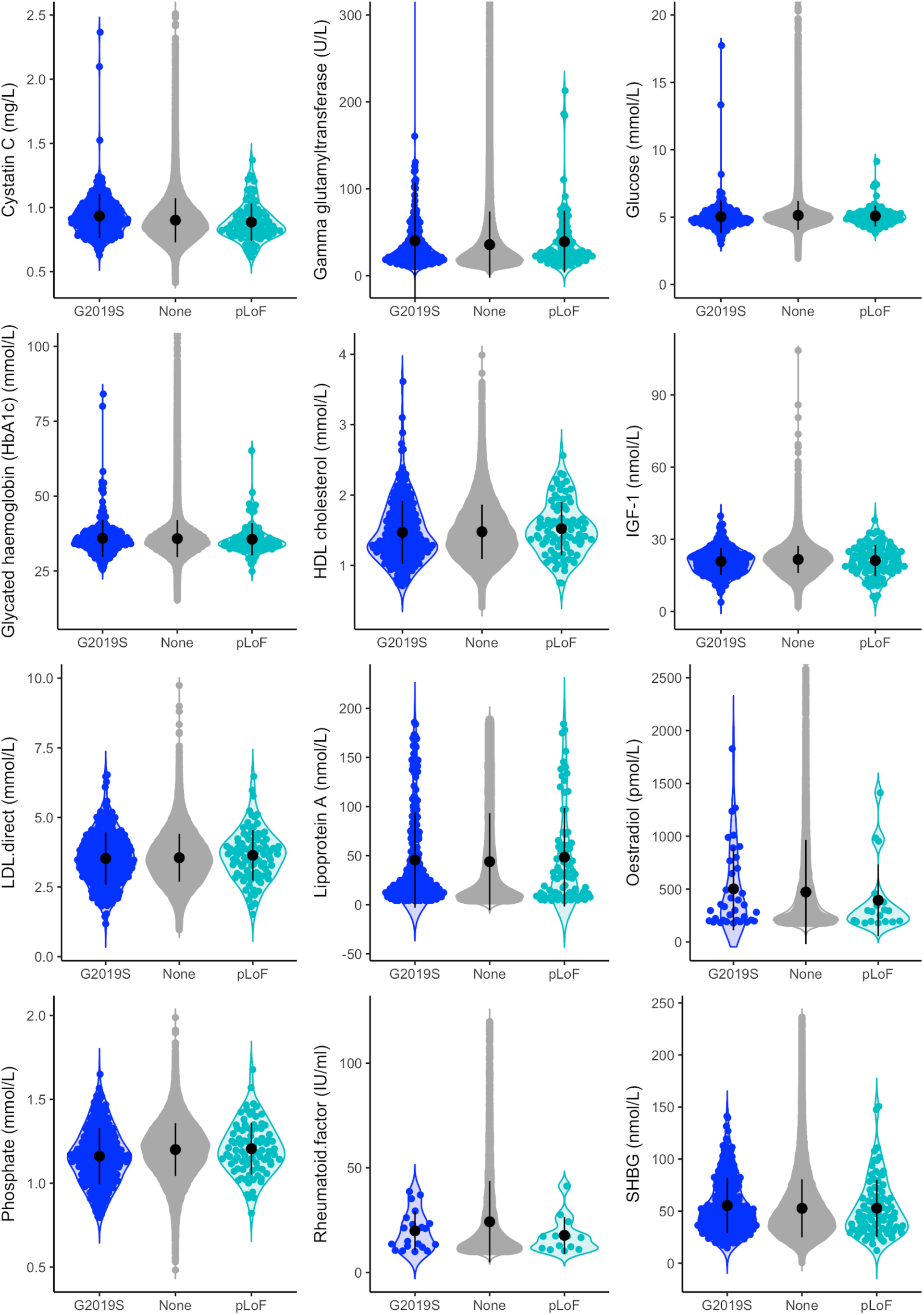
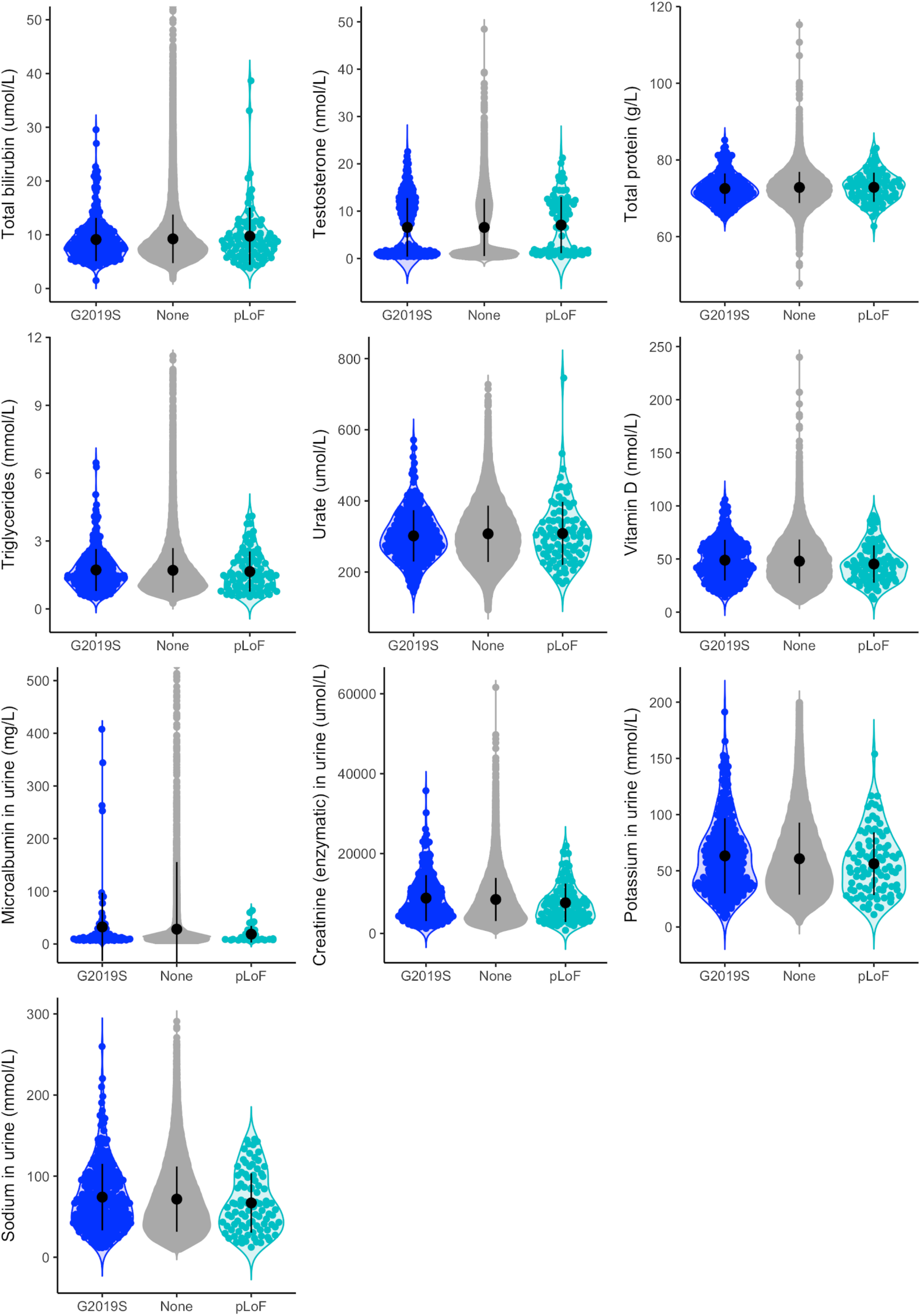
Comparison of 30 blood serum and 4 urine biomarkers between *LRRK2* pLoF carriers (teal), G2019S risk allele carriers (blue) and carriers of neither (None; grey). The mean and standard deviation in each cohort are shown by black circles and lines respectively. For some biomarkers plots have been top truncated to remove outlines in the non-carrier cohort. Values for all pLoF and G2019S carriers are shown within each plot area. In each case, summary statistics are calculated on the full dataset.

**Supplementary Figure 4.**
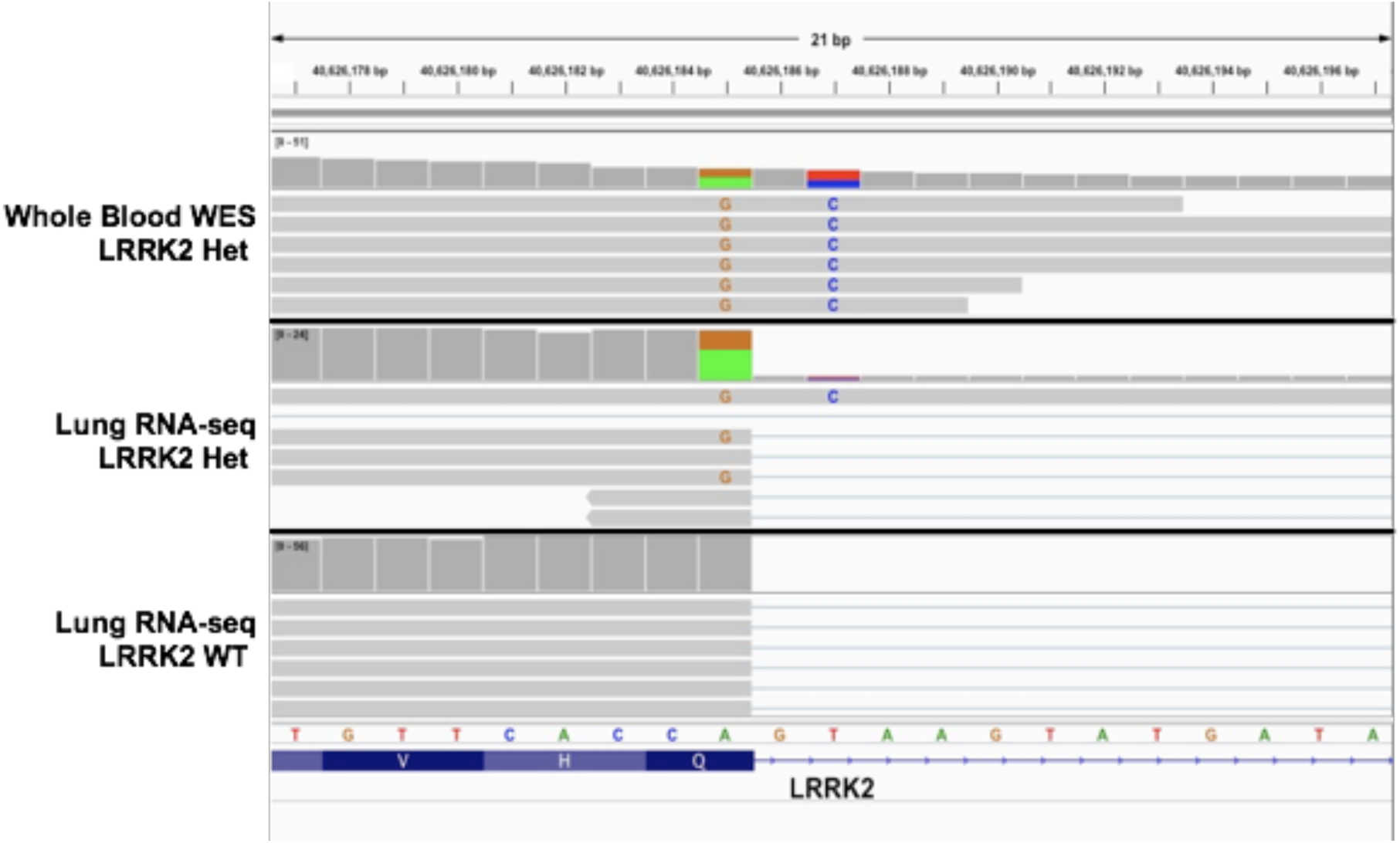
IGV visualization of the splice donor variant GRCh37:12-40626187-T-C in the GTEx *LRRK2* pLoF carrier exome sequencing data and lung tissue RNA-seq data compared to a control GTEx lung RNA-seq sample. The pLoF variant is observed on reads containing an anchoring missense variant, GRCh37:12-40626185-A-G (A green and G orange), and these reads are presenting normal splicing as seen in the control RNA-seq sample.

**Supplementary Figure 5:**
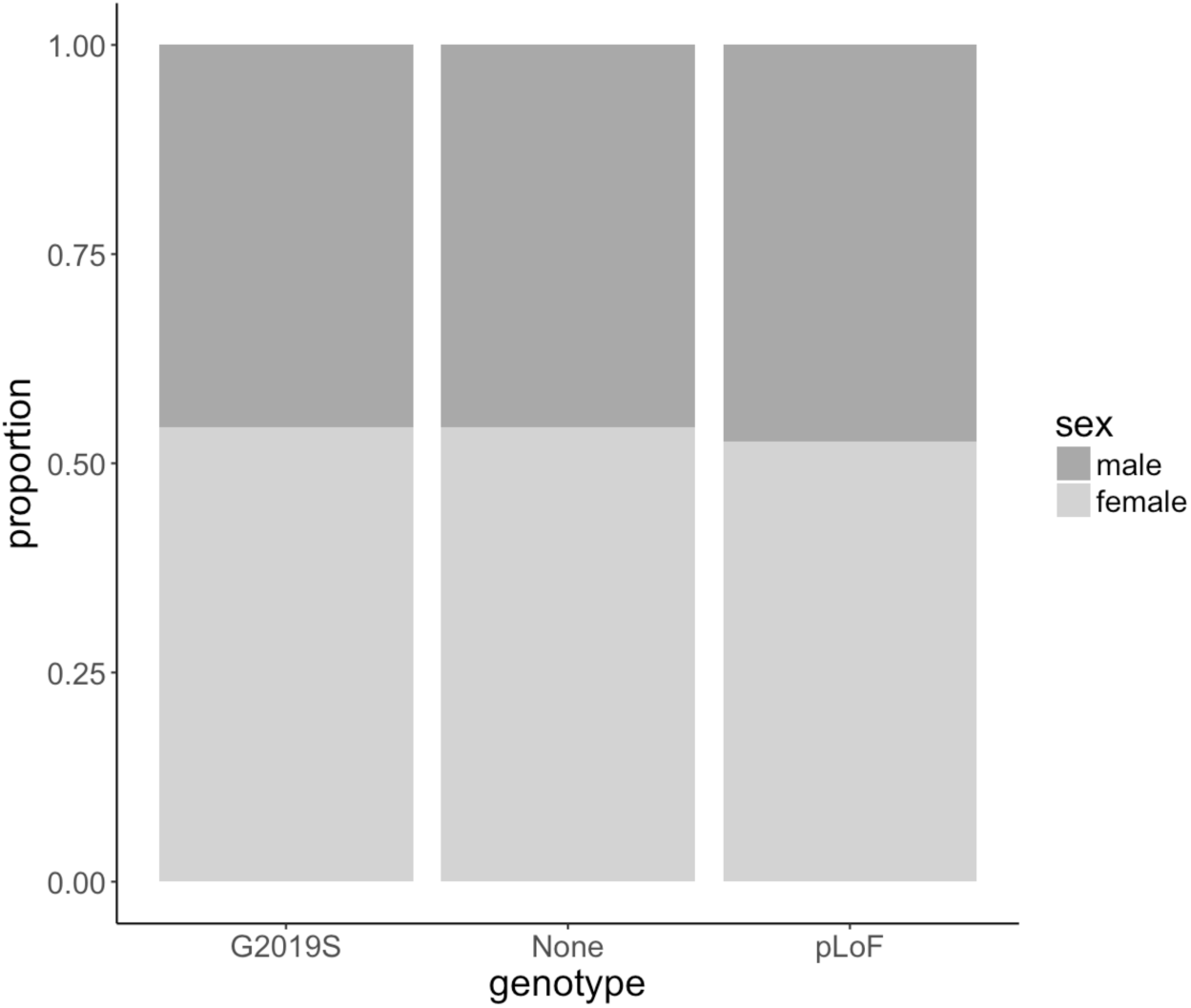
Proportion of *LRRK2* pLoF carriers, G2019S risk allele carriers and non-carriers in the UK Biobank that are male (dark grey) and female (light grey).

**Supplementary Figure 6:**
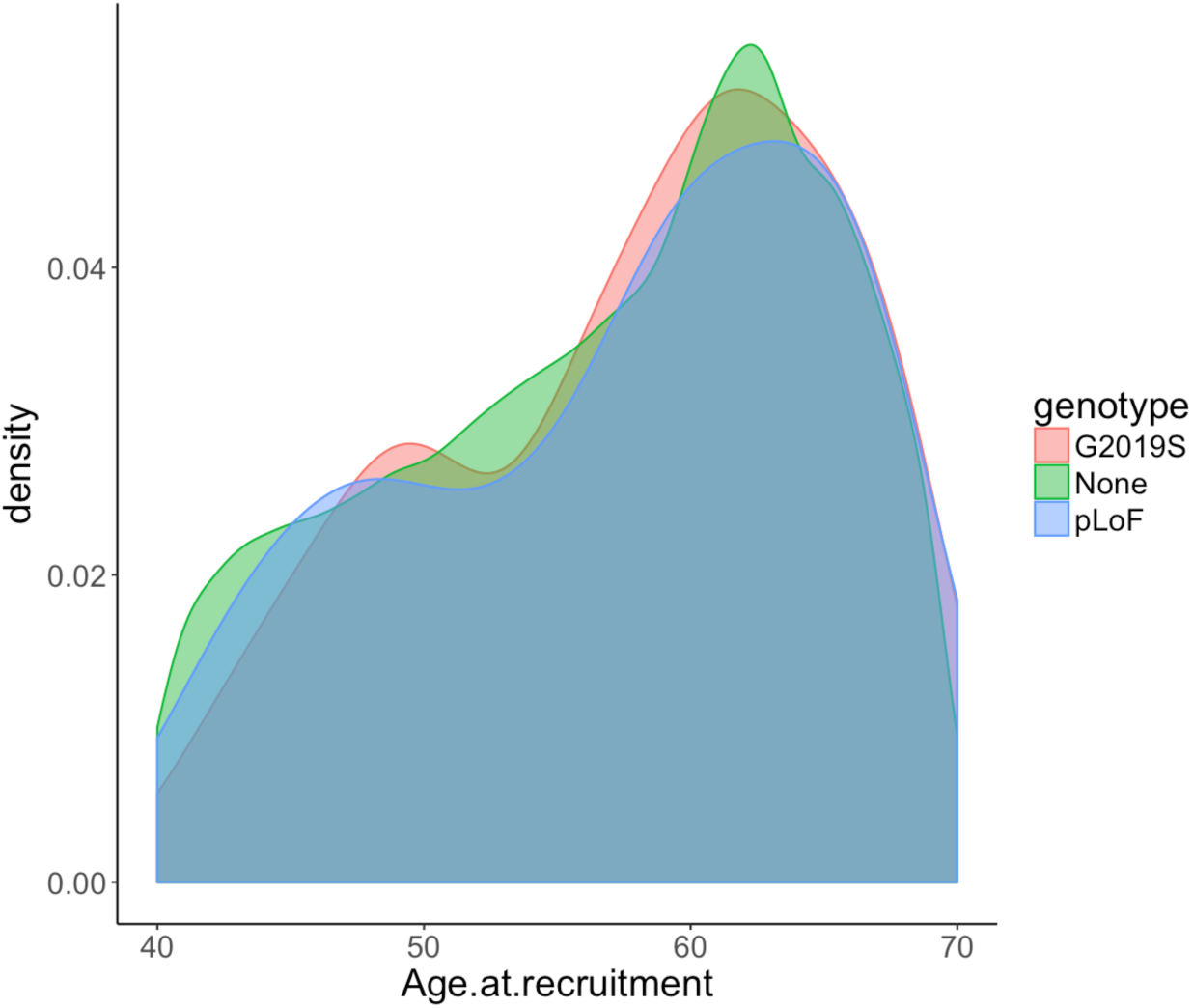
Age distribution of *LRRK2* pLoF carriers, G2019S risk allele carriers and non-carriers in the UK Biobank.

## Online Methods

### gnomAD variant annotation and curation

The gnomAD resource, including both sample and variant quality control (including sample ancestry assignment), is fully described in our companion paper^9^. gnomAD version 2.1.1 was used for this analysis. Putative LoF variants were defined as stop gained, frameshift or essential splice site (splice donor or splice acceptor) as annotated by the Ensembl Variant Effect Predictor^38^.

Variants were included if they were annotated as LoF on any of the three high-confidence GENCODE annotated protein-coding transcripts that are expressed in the lung, liver or kidney. All variants also underwent transcript expression-aware annotation which evaluates the cumulative expression status of the transcripts harboring a variant in the GTEx dataset^39^. All high-confidence variants were found in exons with high evidence of expression across all relevant tissues in GTEx. In addition, all were high-confidence pLoF on the canonical transcript, which is the only transcript to include the kinase domain.

Variants were filtered out if they were flagged as low-confidence by the Loss-of-function Transcript Effect Estimator (LOFTEE)^9^. For the remaining variants, a manual curation was performed, including inspection of variant quality metrics, read distribution and the presence of nearby variants using the Integrative genome viewer (IGV) and splice site prediction algorithms using Alamut.

A single splice-site variant (12-40626187-T-C), found in 77 gnomAD carriers, was identified in an individual with RNAseq data in the Genotype-Tissue Expression (GTEx) project. The RNA-seq reads were manually inspected to look for any effect on splicing. Assessing the read distribution of a linked heterozygous variant in this individual showed convincingly that the variant has no discernible effect on transcript splicing (Supplementary Figure 4). All available tissues were assessed with reads from lung tissue shown in Supplementary Figure 4. The variant was also identified in 8 UK Biobank carriers and in 23andMe and was similarly excluded from these cohorts.

We have complied with all relevant ethical regulations. This study was overseen by the Broad Institute’s Office of Research Subject Protection and the Partners Human Research Committee. Informed consent was obtained from all participants.

### Sanger validation of gnomAD variant carriers

Sanger validation was performed on genomic DNA derived from peripheral blood under the following PCR conditions: 98°C 2min; 30 cycles 20sec 98°C, 20sec 54°C, 1min 72°C; 3min 72°C using the Herculase II Fusion DNA polymerase (Agilent, 600679). PCR products (5ul) were analyzed on a 2% agarose gel and the remaining product was purified with the Qiagen PCR Purification kit. Sequence analysis was performed with both PCR primers at Quintarabio. Details of variants and PCR primers used for each can be found in Supplementary Table 3.

### gnomAD phenotype curation and cohort descriptions

The below described studies with *LRRK2* pLoF carriers had available phenotype data. For each study, all available records were manually reviewed to identify any reports of health problems which were categorised into the following classes: lung, liver, kidney, cardiovascular, nervous system, immune and cancer.

#### The Genomic Psychiatry Cohort (GPC) project

The GPC cohort is a longitudinal resource with the aim of making population-based data available through the National Institute of Mental Health (NIMH). The NIMH repository contains whole-genome sequencing data and detailed clinical and demographic data particularly focussed on schizophrenia and bipolar disorders. A large fraction of participants, 88%, have consented for re-contact^40^.

The screening questionnaire consisted of 32 yes/no questions about mental health issues and 23 yes/no questions covering other medical problems including liver, digestive and cardiovascular problems. There were no specific questions relating to lung or kidney phenotypes although participants were asked to answer yes/no to having any additional health problems. If a participant answered yes to this question, we marked the existence of lung or kidney disease as ‘unknown’. One sample was excluded due to conflicting questionnaire answers.

The age of the 25 *LRRK2* carriers ranged from 19 to 67. Two carriers, aged 55 and 60, reported having had liver problems with 4 participants over 60 years reporting no liver problems.

#### The Pakistan Risk of Myocardial Infarction Study (PROMIS)

The PROMIS cohort comprises 10,503 individuals characterized using a phenotype questionnaire with >350 items covering demographic and dietary characteristics and over 80 blood biomarker measurements^41^. The predominant focus of the questionnaire was cardiac function and phenotype. Whist the participants were specifically asked to report suffering from asthma, no other lung, liver, kidney, nervous system or immune phenotypes were directly assayed so these were marked as ‘unknown’ for these individuals. The 12 *LRRK2* LoF carriers in PROMIS did not differ in terms of age, sex and myocardial infarction status when compared to the entire cohort.

#### The Swedish Schizophrenia and Bipolar Studies

Cases with schizophrenia or bipolar disorder were identified from Swedish national hospitalization registers^42,43^. Controls were selected at random from population registers. All individuals had whole exome sequencing data^44^. All available ICD codes from inpatient hospitalizations and outpatient specialist treatment contacts were provided for each patient.

#### FINRISK: the National FINRISK study

The FINRISK survey has been carried out for 40 years since 1972 every five years using independent, random and representative population samples from different parts of Finland. For this work, we used sequencing and health register data from FINRISK surveys 1992—2007^45^.

Full health records including ICD10 codes were reviewed by study coordinators who provided us with yes/no answers for each of our phenotype classes.

#### The BioMe biobank at The Charles Bronfman Institute for Personalized Medicine at Mount Sinai

The Mount Sinai BioMe Biobank, founded in September 2007, is an ongoing, broadly-consented electronic health record (EHR)-linked bio- and data-repository that enrolls participants non-selectively from the Mount Sinai Medical Center patient population (New York City). BioMe participants represent a broad racial, ethnic and socioeconomic diversity with a distinct and population-specific disease burden, characteristic of the communities served by Mount Sinai Hospital. Currently comprised of over 47,000 participants, BioMe participants are of African (AA, 24%), Hispanic/Latino (HL, 35%), European (EA, 32% of whom 40% Ashkenazi Jewish) and other/mixed ancestry.

BioMe is linked to Mount Sinai’s system-wide Epic EHR, which captures a full spectrum of biomedical phenotypes, including clinical outcomes, covariate and exposure data from past, present and future health care encounters. The median number of outpatient encounters is 21 per participant, reflecting predominant enrollment of participants with common chronic conditions from primary care facilities. Clinical phenotype data has been meticulously harmonized and validated.

Genome-wide genotype data and whole exome sequencing data is available for >30,000 participants. In addition, whole genome sequencing data is available for >11,000 participants. The full electronic records of three BioMe *LRRK2* pLoF carriers were reviewed by local clinicians and we were provided with detailed summaries.

#### Estonian Biobank of the Estonian Genome Center, University of Tartu

The Estonian Biobank cohort is composed of volunteers from the general Estonian resident adult population^46^. The current number of participants - close to 165,000--represents 15%, of the Estonian adult population, making it ideally suited to population-based studies. Participants were recruited throughout Estonia by medical personnel, and participants receive a standardized health examination, donate blood, and fill out a 16-module questionnaire on health-related topics such as lifestyle, diet and clinical diagnoses. A detailed phenotype summary from a health survey and linked data including ICD10 codes, clinical lab values and treatment and medication information is annually updated through linkage with national electronic health databases and registries.

### UK Biobank variant detection and curation

The 49,960 exome sequenced individuals from the UK Biobank were restricted to a subset of 46,062 unrelated individuals of European ancestry. Relatedness was determined using KING kinship coefficient estimates from the genotype relatedness file with a cutoff of 0.0884 to include pairs of individuals with greater than 3rd-degree relatedness. European ancestry was determined by projecting individuals on to 1000 Genomes phase 3^34^ principal component analysis coordinate space, followed by Aberrant R package^47^ clustering to retain only individuals falling within 1000 Genomes project EUR PC1 and PC2 limits (lambda=4.5). We further removed individuals that self-reported as non-European ethnicity.

We identified all individuals with putative LoF variants detected in the FE analysis pipeline using GATK 3.0 for variant calling and filtering^33^. We did not use the SPB pipeline calls due to advertised errors in the Regeneron Genetics Center pipeline at the time we were conducting these analyses. Variants were included if they were annotated as LoF on any transcript expressed in the lung, liver or kidney. As with the gnomAD analysis, variants were filtered out if they were flagged as low-confidence by LOFTEE, before manual curation of the remaining variants. This curation included inspection of variant quality metrics, read distribution and the presence of nearby variants using IGV and splice site prediction algorithms using Alamut.

In addition, 266 individuals in the full genotyped cohort of 488,288 samples who were carriers of the G2019S risk allele were identified. One individual who was a carrier for both a *LRRK2* pLoF variant and G2019S was excluded from all analyses. Carriers of G2019S were not included in the ‘non-carrier’ cohort in any of the analyses.

*LRRK2* pLoF carriers, G2019S risk allele carriers and non-carriers are well matched for both sex (Supplementary Figure 5) and age (Supplementary Figure 6).

### UK Biobank phenotype analysis

#### Blood serum and urine biomarkers

The first recorded value of all fields relating to ‘Blood biochemistry’ (field codes 30600-30890) and ‘Urine assays’ (field codes 30510-30535) was extracted for all individuals. The distribution of values for all biomarkers was plotted (Supplementary Figure 2), and a two-sided Wilcoxon test was used to test for a difference between *LRRK2* pLoF carriers and non-carriers.

These data were also extracted for G2019S risk allele carriers and these individuals were compared to pLoF carriers. There was no significant difference in any of the 34 biomarkers between pLoF and G2019S carriers after accounting for multiple testing (Supplementary Table 14).

#### Clinical measures of kidney function

Albumin to creatinine ratio (ACR) was calculated by dividing the urine microalbumin value (field 30500; mg/L) by the urine creatinine value (field 30510; umol/L) multiplied by a factor of 0.0001131222. Glomerular filtration rate (eGFR) was calculated using the CKD Epidemiology Collaboration (CKD-EPI) creatinine equation^48^. Normal range values for both ACR and eGFR were taken from the National Kidney Foundation website (https://www.kidney.org/kidneydisease/).

#### Spirometry measures of lung function

To assess lung function we used Global Lung Initiative (GLI) 2012 reference equation Z-scores standardised for age, sex and height for Forced expiratory volume in 1-second (FEV1), Forced vital capacity (FVC) and FEV1/FVC ratio measured using Spirometry. These calculations are available in field codes 20256, 20257 and 20258 and were described previously^36^.

#### Grouped phenotype analysis

The list of all codings within the field ‘20002 Non-cancer illness code, self-reported’, were taken from the UK Biobank showcase (http://biobank.ctsu.ox.ac.uk/crystal/coding.cgi?id=6). All selectable codings were given a primary grouping pertaining to the main system relating to that disease. In rare instances where more than one grouping could be assigned, the second was included as a secondary grouping. Diseases with an autoimmune basis were given a secondary grouping to reflect a similar underlying mechanism. Due to the opposing effects of some respiratory diseases, where appropriate, phenotypes in this category were given a secondary grouping of Airway, Interstitial or Pleural. Any codings reflecting symptoms, trauma/injury, benign cancer, mental health phenotypes or diseases arising as a result of infection were excluded. All phenotype codings and assigned groupings are listed in Supplementary Table 9. Any coding within the field ‘20001 Cancer code, self-reported’ was assigned a grouping of ‘Cancer’.

To test for an association between any phenotype group and *LRRK2* pLoF carrier status each individual was counted once as either having self-reported any of the phenotypes within a group, or having reported none. A Fisher’s exact test was used to test for an association.

#### Analysis of ICD10 codes

All ICD10 codes relating to diseases of the liver (K70-K77), diseases of the respiratory system not specific to the upper respiratory tract (J20-J22, J40-J47, J80-J99) or kidney diseases (N00-N29) were curated to exclude any with a primary infectious or external cause (Supplementary Table 11). Asthma was excluded from all analyses to avoid any issues caused by the deliberate ascertainment of the exome sequenced portion of the cohort based on asthma status.

For each individual, we extracted all ICD10 codes from the fields ‘41270 Diagnoses - ICD10’ (recorded from episodes in hospital), ‘40001 Underlying (primary) cause of death: ICD10’ and ‘40002 Contributory (secondary) causes of death: ICD10’. The number of carriers and non-carriers with any ICD10 code relating to lung (5 pLoF carriers; 2,378 non-carriers), liver (0 pLoF carriers; 652 non-carriers) or kidney disease (3 pLoF carriers; 2,272 non-carriers) were counted. For J43 (Emphysema), J44 (Other chronic obstructive pulmonary disease) and J47 (Bronchiectasis), ICD10 codes were not counted if they were reported alongside exposure to or history of tobacco use (Z77.22, P96.81, Z87.891, Z57.31, F17 or Z72.0).

### 23andMe variant annotation and curation

Putative *LRRK2* LoF variants were defined as those classified as high-confidence by LOFTEE. Variants were manually assessed for call rate, genotyping and imputation quality and manually curated to ensure they were expected to cause true LoF.

For each of the two genotyped *LRRK2* pLoF, we determined carrier status by manually inspecting and custom calling the probe intensity plots. For the imputed variants, carrier status was determined from the Minimac-Imputed dosage. As these calling methods might produce false positives, we confirmed the participants’ genotypes through Sanger sequencing. Individuals with discordant genotypes were excluded. This resulted in a cohort of 749 individuals, each of whom is a Sanger sequence-confirmed carrier for one of three pLoF variants (Supplementary Table 4).

During initial selection and sequencing, expansion of the database led to inclusion of a number of additional individuals genotyped for one of the pLoF variants, rs183902574. We performed custom-calling on these individuals and found 354 deemed high-confidence carriers (Supplementary Table 4). As these individuals were not Sanger sequenced, all subsequent analyses were performed both including and excluding these individuals.

Participants provided informed consent and participated in the research online, under a protocol approved by the external AAHRPP-accredited IRB, Ethical & Independent Review Services (E&I Review).

### Testing the power to detect an age effect in 23andMe

As a positive control for the age analysis, we tested the APOE Alzheimer’s disease risk allele rs429358, which has a known effect on lifespan. This effect is highly significant in this dataset (*P*=1.2×10^−211^).

Given that the carrier count for rs429358 is much higher than for *LRRK2* pLoF, we assessed the power of the 23andMe dataset to detect an age effect associated with LRRK2 pLoF variants that is of the same effect size as the known effect of the APOE allele rs429358 by sampling carriers of this variant. We randomly selected N carriers of rs429358 from the 23andMe dataset, performed a KS test on the age distribution of those carriers versus 4,000,000 non-carriers, and considered the resulting p-value. We repeated this process 100 times, and then computed the proportion of these simulations with p < 0.05. This tells us our power to reject the null hypothesis that APOE does not have an affect on age at alpha=0.05, if we had N carriers in the dataset. We repeated this for different values of N, between 1,000 and 20,000 (Supplementary Table 15).

### Association testing in the 23andMe dataset

#### Phenotype selection

The 23andMe dataset includes self-reported phenotype data for thousands of phenotypes across a diverse range of categories. These phenotypes have different sample sizes and prevalences, so the power to detect associations varies widely. We began with a curated set of 748 disease phenotypes. We then applied a liberal filter based on our power to detect an association with carrier status. More specifically, assuming a MAF of 2E-05, we restricted to phenotypes where we had power 0.1 to detect an association effect with OR > 1.3 (for binary traits) or beta > 0.2 (for quantitative traits) at alpha=0.0001 significance. This left us with 460 binary and 14 quantitative phenotypes.

#### Association testing

For the subset of 368 health-related phenotypes (i.e. excluding any related to diet, drug use, lifestyle and personality), we first restricted to individuals for whom we had phenotypic data. We calculated pairwise identity by descent (IBD) over all individuals using a modified version of IBD64, and then iteratively removed individuals until we were left with a set of participants, no two of whom shared > 700cM in IBD. We then tested the association between the phenotype and carrier status, controlling for age, sex, genotyping platform, and the first ten genetic principle components. We used logistic regression for binary phenotypes, and linear regression for quantitative phenotypes.

To control for population structure we restricted our analyses to participants with > 97% European ancestry, but the results did not qualitatively change when we dropped this restriction. We also tested associations using only individuals whose carrier status was confirmed by Sanger sequencing, but this also did not result in any meaningful difference.

A Bonferroni corrected *P*-value threshold for 368 independent tests of 1.36×10^−4^ was used to assess statistical significance.

#### Power analysis

For each phenotype, we compute the theoretical odds ratio we are powered to detect (given in Supplementary Tables 6 and 7) as follows: Let m be the proportion of individuals used in the association study of that phenotype who are LRRK2 pLoF carriers, and let n0 and n1 be the number of controls and cases, respectively. For each OR in the interval [1, 10] at steps of 0.02, we compute the power of the Cochran-Armitage trend test to detect an association between a variant with MAF m and odds ratio OR at alpha=0.05, with n0 controls and n1 cases^49^. We report the smallest OR such that this power is >= 0.8.

### Analysis of LRRK2 protein levels

#### Cell culture

All cell lines tested negative for mycoplasma contamination on a monthly basis with the MycoAlert™ Detection kit (Lonza, LT07-118) and MycoAlert™ Assay Control Set (Lonza, LT07-518). Cells were grown at 37°C with 5% CO2.

#### Human embryonic stem cell culture

All pluripotent stem cells were approved by Harvard ESCRO protocol #E00052 and #E00067. Human embryonic stem cells (hESCs) were obtained from WiCell Research Institute (WA01, H1) under an MTA. Cell lines were authenticated by visual inspection of cell morphology with brightfield microscopy, staining with anti-Oct4 antibody to determine maintenance of pluripotency (Santa Cruz, sc-5279, data not shown), sent to WiCell Research Institute after 6 months of passaging or after isogenic cell line generation for karyotyping, and in some cases PCA of RNA sequencing data to confirm clustering with other pluripotent stem cell lines. Pluripotent stem cells were plated onto hESC qualified matrigel (VWR, BD354277) coated 6 well plates, mTeSR1 media was changed daily (StemCell Technologies, 85850), and cells were passaged every 5-7 days with 0.5mM EDTA.

#### Lymphoblastoid cell culture

Lymphoblastoid cell lines (LCLs) were obtained from Coriell Biorepository (GM18500, GM18501, GM18502, HG01345, HG01346) and approved by the Broad Institute Office of Research Subject Protection protocol #3639. Cell lines were authenticated by visual inspection of cell morphology with brightfield microscopy and in some cases PCA of RNA sequencing data to confirm clustering with GTEx LCLs. LCL media was changed every other day with RPMI 1640 (Life Technologies), 2mM L-glutamine (Life Technologies), and 15% FBS (Sigma).

#### Cardiomyocyte differentiation

Cardiomyocyte differentiation of the control and engineered H1 hESC lines was performed according to the protocol by Lian et al., 2012. Briefly, 500,000 cells were plated on hESC qualified matrigel (VWR, BD354277), grown in mTeSR1 media for four days (StemCell Technologies, 85850), and switched to RPMI media (Life Technologies) with B27 supplement (Life Technologies), switching to B27 with insulin at day 7, for the remainder of the protocol. On day 0 of differentiation, 12µM CHIR99021 (Tocris) was applied for 24 hours. At day 3, cultures were treated with 5µM IWP2 (Tocris) for 24 hours. Bright field images and movies were acquired at day 17 and cells were harvested for protein/RNA extraction at day 19.

#### Isogenic cell line engineering

The following guide and HR template were delivered into single cell H1 hESCs via nucleofection (Lonza 4D-Nucleofector X unit) using the P3 Primary Cell Kit (V4XP-3024), pulse code CA137, and pX459 (Addgene): AATAAGGCATTTCATATAGT and ACAGGCCTGTGATAGAGCTTCCCCATTGTGAGAACTCTGAAATTATCATCTGACTATATGAAATGCCTTATTTTCCAATGGGATTTTGGTCAAGATTAA. Cells were allowed to recover from nucleofection in mTeSR supplemented with 10µM Rock Inhibitor (Y-27632, Tocris) overnight. The following three days the cells were treated with 0.25µg/ml puromycin (VWR) in mTeSR. Cells were then cultured in mTeSR until colonies were ready to be split. Engineered cells were split into single cells and plated in matrigel coated 96-well plates at a density of 0.5 cells per well. Plates were screened for colonies 8-10 days after plating and grown until colonies were ready to be split. Colonies were then split with 0.5mM EDTA into two identical 96-well plates, one for DNA extraction/PCR/sequencing and one for freezing cells. Once colonies were ready to be split, 96-well plates were frozen in mFreSR (Stem Cell Technologies) and stored at −80°C until HR positive wells were identified. HR edited cells were then thawed and expanded for 4 generations, validated by Sanger sequencing, karyotyping, and OCT4 staining prior to proceeding with cardiomyocyte differentiation.

#### Off-target analysis of CRISPR/Cas9 engineering

To detect any potential off-target effects caused by CRISPR/Cas9 genome editing, whole-genome sequencing (WGS) was conducted for both the engineered and control cell lines. DNA extraction, Quality Control (QC) and 30x PCR-Free WGS were performed by the Genomics Platform at the Broad Institute. AllPrep DNA/RNA extraction kit was used, following its protocol. Alignment, marking of duplicates, recalibration of quality scores and variant calling are all performed using GATK best practices^50^.

We identified 157,230 variants in the engineered cell line that were not found in the control cell line as candidate variants. For the guide RNA (gRNA) used, we defined potential off-target regions as those with <4bps mismatch against the full 20bp gRNA sequence (334 regions), and/or <2bp mismatch against the seed (=PAM proximal) 12bp of the gRNA sequence (5,780 regions), each followed by the NGG PAM. We looked for any candidate variant that fall into the potential off-target region, resulting in detection of only one variant (chr8-65084564-A-AT) that falls onto a region with 1 mismatch against the seed 12bp of gRNA sequence (chr8:65084560-65084575). No variants with <4bp mismatch against the full 20bp gRNA sequence or perfect match at the seed region were detected. Since a mismatch at the seed region decreases the likelihood of off-target variants^51,52^, and also because the single variant we detected is a known variant (rs1161563412) observed in the population without apparent phenotypic association, we concluded no major off-target effect exists at the level of violating the main steps of our research. All the analysis for the detection of potential off-target were conducted using pybedtools^53^ and CRISPRdirect^54^ software.

#### Western blot analysis

Cell pellets were snap-frozen in liquid nitrogen and stored at −80°C. Cells were dounce homogenized in ice-cold radioimmunoprecipitation assay buffer (RIPA) (89901; Thermo Scientific) containing protease inhibitors (HaltTM Protease Inhibitor, Thermo Scientific). Homogenates were rotated at 4°C for 30min, followed by centrifugation at 15,000g for 20min at 4°C. Equal amounts of protein (50µg) were electrophoresed on 4-20% SDS-PAGE (Biorad) and transferred to nitrocellulose membranes. The following antibodies were used for immunoblotting: LRRK2 (75-253, UC Davis/NIH NeuroMab Facility), α-actinin (A7811, Sigma), GAPDH (sc-25778, Santa Cruz), anti-rabbit IgG HRP (7074, Cell Signaling), and anti-mouse IgG HRP (7076, Cell Signaling). Immunoblots were developed using enhanced chemiluminescence (SuperSignal West Pico Chemiluminescent Substrate, Thermo Scientific) on an Amersham Imager 600.

## Notes

#### Summary of Updates

Addition of the UK Biobank cohort

## References

1. Nelson, M. R. et al. The support of human genetic evidence for approved drug indications. Nat. Genet. 47, 856–860 (2015).

2. Plenge, R. M., Scolnick, E. M. & Altshuler, D. Validating therapeutic targets through human genetics. Nat. Rev. Drug Discov. 12, 581–594 (2013).

3. Greggio, E. et al. Kinase activity is required for the toxic effects of mutant LRRK2/dardarin. Neurobiol. Dis. 23, 329–341 (2006).

4. West, A. B. et al. Parkinson’s disease-associated mutations in leucine-rich repeat kinase 2 augment kinase activity. Proc. Natl. Acad. Sci. U. S. A. 102, 16842–16847 (2005).

5. Andersen, M. A. et al. PFE-360-induced LRRK2 inhibition induces reversible, non-adverse renal changes in rats. Toxicology 395, 15–22 (2018).

6. Fuji, R. N. et al. Effect of selective LRRK2 kinase inhibition on nonhuman primate lung. Sci. Transl. Med. 7, 9273ra15 (2015).

7. Baptista, M. A. S. et al. Loss of leucine-rich repeat kinase 2 (LRRK2) in rats leads to progressive abnormal phenotypes in peripheral organs. PLoS One 8, e80705 (2013).

8. Hinkle, K. M. et al. LRRK2 knockout mice have an intact dopaminergic system but display alterations in exploratory and motor co-ordination behaviors. Mol. Neurodegener. 7, 25 (2012).

9. Karczewski, K. J. et al. Variation across 141,456 human exomes and genomes reveals the spectrum of loss-of-function intolerance across human protein-coding genes. bioRxiv 531210 (2019). doi:10.1101/531210

10. de Lau, L. M. L. & Breteler, M. M. B. Epidemiology of Parkinson’s disease. Lancet Neurol. 5, 1. 525–535 (2006).

11. Polymeropoulos, M. H. et al. Mapping of a gene for Parkinson’s disease to chromosome 4q21-q23. Science 274, 1197–1199 (1996).

12. Klein, C. & Westenberger, A. Genetics of Parkinson’s disease. Cold Spring Harb. Perspect. Med. 2, a008888 (2012).

13. Zimprich, A. et al. Mutations in LRRK2 cause autosomal-dominant parkinsonism with pleomorphic pathology. Neuron 44, 601–607 (2004).

14. Goldwurm, S. et al. Evaluation of LRRK2 G2019S penetrance: relevance for genetic counseling in Parkinson disease. Neurology 68, 1141–1143 (2007).

15. Do, C. B. et al. Web-based genome-wide association study identifies two novel loci and a substantial genetic component for Parkinson’s disease. PLoS Genet. 7, e1002141 (2011).

16. MacLeod, D. et al. The familial Parkinsonism gene LRRK2 regulates neurite process morphology. Neuron 52, 587–593 (2006).

17. West, A. B. et al. Parkinson’s disease-associated mutations in LRRK2 link enhanced GTP-binding and kinase activities to neuronal toxicity. Hum. Mol. Genet. 16, 223–232 (2007).

18. Steger, M. et al. Phosphoproteomics reveals that Parkinson’s disease kinase LRRK2 regulates a subset of Rab GTPases. Elife 5, (2016).

19. Roosen, D. A. & Cookson, M. R. LRRK2 at the interface of autophagosomes, endosomes and lysosomes. Mol. Neurodegener. 11, 73 (2016).

20. Di Maio, R. et al. LRRK2 activation in idiopathic Parkinson’s disease. Sci. Transl. Med. 10, (2018).

21. Zhao, H. T. et al. LRRK2 Antisense Oligonucleotides Ameliorate a-Synuclein Inclusion Formation in a Parkinson’s Disease Mouse Model. Mol. Ther. Nucleic Acids 8, 508–519 (2017).

22. Chen, Z.-C.. et al. Phosphorylation of amyloid precursor protein by mutant LRRK2 promotes AICD activity and neurotoxicity in Parkinson’s disease. Sci. Signal. 10, (2017).

23. Chen, J., Chen, Y. & Pu, J. Leucine-Rich Repeat Kinase 2 in Parkinson’s Disease: Updated from Pathogenesis to Potential Therapeutic Target. Eur. Neurol. 79, 256–265 (2018).

24. Daniel, G. & Moore, D. J. Modeling LRRK2 Pathobiology in Parkinson’s Disease: From Yeast to Rodents. in Behavioral Neurobiology of Huntington’s Disease and Parkinson’s Disease (eds. Nguyen, H. H. P. & Cenci, M. A.) 331–368 (Springer Berlin Heidelberg, 2015).

25. Cohen, J. C., Boerwinkle, E., Mosley, T. H., Jr & Hobbs, H. H. Sequence variations in PCSK9, low LDL, and protection against coronary heart disease. N. Engl. J. Med. 354, 1264–1272 (2006).

26. TG and HDL Working Group of the Exome Sequencing Project, National Heart, Lung, and Blood Institute et al. Loss-of-function mutations in APOC3, triglycerides, and coronary disease. N. Engl. J. Med. 371, 22–31 (2014).

27. Myocardial Infarction Genetics Consortium Investigators et al. Inactivating mutations in NPC1L1 and protection from coronary heart disease. N. Engl. J. Med. 371, 2072–2082 (2014).

28. Minikel, E. V. et al. Quantifying prion disease penetrance using large population control cohorts. Sci. Transl. Med. 8, 322ra9 (2016).

29. Lek, M. et al. Analysis of protein-coding genetic variation in 60,706 humans. Nature 536, 285–291 (2016).

30. MacArthur, D. G. et al. A systematic survey of loss-of-function variants in human protein-coding genes. Science 335, 823–828 (2012).

31. Minikel, E. V. et al. Evaluating potential drug targets through human loss-of-function genetic variation. bioRxiv 530881 (2019). doi:10.1101/530881

32. Blauwendraat, C. et al. Frequency of Loss of Function Variants in LRRK2 in Parkinson Disease. JAMA Neurol. 75, 1416–1422 (2018).

33. Van Hout, C. V. et al. Whole exome sequencing and characterization of coding variation in 49,960 individuals in the UK Biobank. bioRxiv 572347 (2019). doi:10.1101/572347

34. 1000 Genomes Project Consortium et al. A global reference for human genetic variation. Nature 526, 68–74 (2015).

35. UK10K Consortium et al. The UK10K project identifies rare variants in health and disease. Nature 526, 82–90 (2015).

36. Gupta, R. P. & Strachan, D. P. Ventilatory function as a predictor of mortality in lifelong non-smokers: evidence from large British cohort studies. BMJ Open 7, e015381 (2017).

37. Troyer, M. Safety, Tolerability and Target Engagement Demonstrated in Phase 1 Study of LRRK2 Inhibitor DNL201 in Healthy Young and Elderly Adults. in https://denalitherapeutics.com/uploads/documents/events/DNL201_MJFF_PDTx_Presentation.pdf (MJFF Parkinson’s Disease Therapeutics Conference, 2018).

38. McLaren, W. et al. The Ensembl Variant Effect Predictor. Genome Biol. 17, 122 (2016).

39. Cummings, B. B. et al. Transcript expression-aware annotation improves rare variant discovery and interpretation. bioRxiv 554444 (2019). doi:10.1101/554444

40. Pato, M. T. et al. The genomic psychiatry cohort: partners in discovery. Am. J. Med. Genet. B Neuropsychiatr. Genet. 162B, 306–312 (2013).

41. Saleheen, D. et al. Human knockouts and phenotypic analysis in a cohort with a high rate of consanguinity. Nature 544, 235–239 (2017).

42. Ripke, S. et al. Genome-wide association analysis identifies 13 new risk loci for schizophrenia. Nat. Genet. 45, 1150–1159 (2013).

43. Bipolar Disorder and Schizophrenia Working Group of the Psychiatric Genomics Consortium. Electronic address: & Bipolar Disorder and Schizophrenia Working Group of the Psychiatric Genomics Consortium. Genomic Dissection of Bipolar Disorder and Schizophrenia, Including 28 Subphenotypes. Cell 173, 1705–1715.e16 (2018).

44. Genovese, G. et al. Increased burden of ultra-rare protein-altering variants among 4,877 individuals with schizophrenia. Nat. Neurosci. 19, 1433–1441 (2016).

45. Borodulin, K. et al. Cohort Profile: The National FINRISK Study. Int. J. Epidemiol. (2017). doi:10.1093/ije/dyx239

46. Leitsalu, L. et al. Cohort Profile: Estonian Biobank of the Estonian Genome Center, University of Tartu. Int. J. Epidemiol. 44, 1137–1147 (2015).

47. Bellenguez, C. et al. A robust clustering algorithm for identifying problematic samples in genome-wide association studies. Bioinformatics 28, 134–135 (2012).

48. Levey, A. S. & Stevens, L. A. Estimating GFR using the CKD Epidemiology Collaboration (CKD-EPI) creatinine equation: more accurate GFR estimates, lower CKD prevalence estimates, and better risk predictions. American journal of kidney diseases: the official journal of the National Kidney Foundation 55, 622–627 (2010).

49. Freidlin, B., Zheng, G., Li, Z. & Gastwirth, J. L. Trend tests for case-control studies of genetic markers: power, sample size and robustness. Hum. Hered. 53, 146–152 (2002).

50. McKenna, A. et al. The Genome Analysis Toolkit: a MapReduce framework for analyzing next-generation DNA sequencing data. Genome Res. 20, 1297–1303 (2010).

51. Hsu, P. D. et al. DNA targeting specificity of RNA-guided Cas9 nucleases. Nat. Biotechnol. 31, 827–832 (2013).

52. Wang, Q. & Ui-Tei, K. Computational Prediction of CRISPR/Cas9 Target Sites Reveals Potential Off-Target Risks in Human and Mouse. in Genome Editing in Animals: Methods and Protocols (ed. Hatada, I.) 43–53 (Springer New York, 2017).

53. Dale, R. K., Pedersen, B. S. & Quinlan, A. R. Pybedtools: a flexible Python library for manipulating genomic datasets and annotations. Bioinformatics 27, 3423–3424 (2011).

54. Naito, Y., Hino, K., Bono, H. & Ui-Tei, K. CRISPRdirect: software for designing CRISPR/Cas guide RNA with reduced off-target sites. Bioinformatics 31, 1120–1123 (2015).

